# Hotspot mutations and ColE1 plasmids contribute to the fitness of *Salmonella* Heidelberg in poultry litter

**DOI:** 10.1101/312322

**Authors:** Adelumola Oladeinde, Kimberly Cook, Alex Orlek, Greg Zock, Kyler Herrington, Nelson Cox, Jodie Plumblee Lawrence, Carolina Hall

## Abstract

*Salmonella enterica* subsp. *enterica* serovar Heidelberg (*S.* Heidelberg) is a clinically-important serovar linked to food-borne illness, and commonly isolated from poultry. Investigations of a large, multistate outbreak in the USA in 2013 identified poultry litter (PL) as an important extra-intestinal environment that may have selected for specific *S.* Heidelberg strains. Poultry litter is a mixture of bedding materials and chicken excreta that contains chicken gastrointestinal (GI) bacteria, undigested feed, feathers, and other materials of chicken origin. In this study, we performed a series of controlled laboratory experiments which assessed the microevolution of two S. Heidelberg strains (SH-2813 and SH-116) in PL previously used to raise 3 flocks of broiler chickens. The strains are closely related at the chromosome level, differing from the reference genome by 109 and 89 single nucleotide polymorphisms/InDels, respectively. Whole genome sequencing was performed on 86 isolates recovered after 0, 1, 7 and 14 days of microevolution in PL. Only strains carrying an IncX1 (37kb), 2 ColE1 (4 and 6kb) and 1 ColpVC (2kb) plasmids survived more than 7 days in PL. Competition experiments showed that carriage of these plasmids was associated with increased fitness. This increased fitness was associated with an increased copy number of IncX1 and ColE1 plasmids. Further, all Col plasmid-bearing strains had hotspot mutations in 37 loci on the chromosome and in 3 loci on the IncX1 plasmid. Additionally, we observed a decrease in susceptibility to tobramycin, kanamycin, gentamicin, neomycin and fosfomycin for Col plasmid-bearing strains. Our study demonstrates how positive selection from poultry litter can change the evolutionary path of S. Heidelberg.

## INTRODUCTION

The United States (U.S.) is the world’s largest producer of poultry with over 8 billion broilers produced yearly (1). Maintaining a safe supply of poultry is increasingly important as demand increases, production becomes more intensive and facilities become antibiotic free. Broilers (around 22,000 broilers per house) are typically raised in open plan barns on a deep floor of bedding materials (typically wood shavings, rice hulls or saw dust (1, 2). Poultry litter (PL) is a mixture of bedding materials and chicken excreta that contains chicken gastrointestinal (GI) bacteria, undigested feed, uric acid, feathers and other materials of chicken origin (3–6). As the cost of bedding materials increase, land nutrient management requirements become more stringent, and broiler production and turnover times decrease, farmers have chosen to increase the number of flocks raised on litter before cleanout. This practice is accepted in production environments, but the effect of litter reuse on microbial populations has not been well studied.

Available data suggest that litter re-use can result in significant changes in the microbiological, chemical and physical composition of litter (3, 7, 8). In addition, reused litter has been found to harbor fewer yeasts, molds, coliform bacteria and *Salmonella* compared to fresh litter (9, 10). However, fecal shedding of food borne pathogens such as *Salmonella* into poultry by colonized birds can also be a source of new infection or re-inoculation. For instance, *Salmonella* in 131-day-old litter was shown to colonize 82% of day-old chicks after placement for 32 days (11) and *Salmonella enterica* subsp. *enterica* serovar Heidelberg (S. Heidelberg) has been shown to persist in recycled litter, even after litter treatment (12). *Salmonella* Heidelberg is a clinically-important serovar, linked to food-borne illness and commonly isolated from poultry. After *S*. Typhimurium, *S*. Heidelberg is the serovar of *Salmonella* most often associated with *Salmonella-related* deaths in the United States (13). In 2013, S. Heidelberg was responsible for a large multistate foodborne outbreak (634 cases) linked to consumption of chicken (14). Epidemiological and environmental investigations identified PL as an important extra-intestinal environment that may have selected for specific *S.* Heidelberg strains (14).

*Salmonella* Heidelberg is highly clonal and cannot be subtyped by conventional pulsed-field gel electrophoresis (PFGE) which poses an inherent difficulty when differentiating strains responsible for foodborne outbreaks from those associated with background sporadic cases (15–17). High quality core genome single nucleotide variant (hqSNV) analysis on whole genome DNA sequences (WGS) of S. Heidelberg has been recommended as a better approach to distinguish outbreak strains (15). However, core genome SNV is non-discriminatory between isolates harboring horizontally acquired antibiotic resistance genes (ARG) versus susceptible ones. Multidrug resistant *S*. Heidelberg isolates tend to cluster with susceptible isolates during outbreaks and differ by only 0 - 4 SNVs (16, 17). In a study by Edirmanasinghe et al. (17), the authors reported 65 out of 113 sequenced *S*. Heidelberg encoded the AmpC beta-lactamase (blaCMY-2) gene on an incompatibility group I1 plasmid (IncI1) and were resistant to important 3^rd^ generation cephalosporins. On the other hand, susceptible isolates (n=38) carried multiple variants of Col plasmids and encoded no known ARG. The function of these Col plasmids is not clear but they seem to be mobilized by co-resident conjugal plasmids, such as IncI1 and IncX, two plasmids frequently found among S. Heidelberg isolates (16–19).

The majority of S. Heidelberg isolates sequenced and available in the National Center for Biotechnology Information (NCBI) database were recovered from either food or clinical samples (18, 20–22), leaving a knowledge gap in our understanding of strains of environmental origin. The poultry litter microbiome may carry mobile genetic elements (MGE) including plasmids that can be horizontally transferred to *S*. Heidelberg upon introduction into PL. These newly acquired MGE may present a fitness advantage to permissive recipients, further enabling the persistence of *S*. Heidelberg in poultry environments.

In this study, we investigated the role of PL on the fitness of two S. Heidelberg strains with indistinguishable PFGE patterns, both recovered from chicken carcass in two states in USA. Both strains harbor an IncX1 plasmid (37 kb) but differ in the occurrence of Col plasmids. We performed a series of controlled laboratory experiments to assess the microevolution of these strains in PL that was previously used to raise three flocks of broiler chickens. Our data suggests that acquisition of multiple ColE1 plasmids in tandem with mutations in *S*. Heidelberg chromosome and the IncX1 plasmid may have contributed to the persistence of *S.* Heidelberg in the PL used in this study. Additionally, the presence of Col plasmids was associated with a decreased susceptibility to tobramycin, kanamycin, neomycin and fosfomycin.

## MATERIALS AND METHODS

### Poultry Litter for microcosm evolution studies

Poultry litter (PL) for microcosm studies was provided by the University of Georgia poultry research center in Athens, GA, USA. The PL was previously used to raise 3 flocks of broiler chickens under simulated commercial poultry production conditions. The litter was collected approximately 2 weeks after the third flocks of birds were removed. The PL bedding material was composed of pine shavings.

### Bacterial strains and preconditioning

S. Heidelberg strains (SH-2813 and SH-116) used in this study were originally recovered from broiler chicken carcass rinsates. The genotypic/phenotypic characteristics of these strains are provided in Table 1. The strains were preconditioned in poultry litter extract (PLE) media prepared as described in supplementary material or in Brain Heart Infusion broth (BHIB; BD Difco, Sparks, MD) before inoculation into PL. For each strain, 5 single colonies from Sheep Blood Agar (SBA) (Remel Inc, San Diego, CA) were transferred to 10 ml of PLE and BHIB, and incubated at 37 ^o^C for 24 h. Serial batch culturing was carried out for approximately 50 generations in PLE and BHIB by transferring 100 μl of culture to 10 ml of fresh PLE and BHIB (100-fold dilution) to yield approximately 10 gen. d^-1^. After 5 days, the population was used as inoculum for evolution experiments in PL microcosms.

**Table 1.** Characteristics of the strains used in this study.

| Strain ID | Serotype | Plasmid Profile (size) | *Resistance profile | #Antimicrobial resistance genes | ¶Unique Single Nucleotide Variants (SNV) |
| --- | --- | --- | --- | --- | --- |
| SH-2813 | ST15 | IncX1 (37kb) | Ery, Tyl, | AAC (6')-Iy, FosA7 | 109 |
| SH-116 | ST15 | IncX1 (37 kb), ColE1(6 kb) ColE1 (4 kb) ColpVC (2 kb) | Ery, Tyl, Pen | AAC (6')-Iy, FosA7 | 89 |

Note. - Ery = Erythromycin; Pen = Penicillin; Tyl= Tylosin
\* Susceptible to all 14 antibiotics on the NARMs Gram-negative AST panel. Ery, Pen and Tyl are on the NARMs Gram-positive AST panel.

### Experimental microevolution in PL

Poultry litter was collected in a 5-gallon bucket and thoroughly mixed before 30 g aliquots were placed into 48 250 ml Nalgene wide jars and 1X PBS (3-4 ml) was added to achieve 30 *%* moisture. Microcosms were incubated for 72 h at 37°C before being individually inoculated with approximately 10^6^ colonies forming units (CFU) of either SH-2813 or SH-116 preconditioned previously in PLE or BHIB. These 4 groups/treatments were as follows: SH-2813 + BHIB, SH-2813 + PLE, SH-116 +BHIB and SH-116 +PLE. Inoculated microcosms were thoroughly mixed and then incubated for 14 days at 37 ^o^C (Supplementary Fig. 1). A control microcosm remained un-inoculated. Three microcosms per group were destructively sampled at days 0, 1, 7 and 14.

### Poultry litter microcosm sampling

Poultry litter samples (25 g) were mixed with 150 ml 1X PBS and 5-10 ml glass beads (Fisher Sci: S800242) then homogenized for 10 min at 450 rpm in a hand wrist shaker (Boekel Scientific, Feasterville, PA). The PL mixtures were allowed to settle for 5 min to allow settling of large debris/pine shavings and then passed through cheesecloth and hand pressed to recover eluates which were used for downstream microbiological and chemical analysis. The moisture content of each PL sample was determined gravimetrically by drying approximately 1 g of PL at 107 °C for 24 h.

### Microbiological methods

Recovered PL eluate was serially diluted in 1X PBS and plated on selective or general-purpose growth media. We used Brilliant Green Sulfa (BGS) and XLT-4 agar for *Salmonella* spp., Chromagar ECC for coliforms, Enterococcosel for *Enterococcus* spp. and Plate Count Agar (PCA) for total bacteria. For enrichment and recovery of injured/viable *Salmonella* spp., 1 ml of eluate was transferred to 20 ml of 1 *%* Buffered Peptone Water (BPW) and incubated 24 h before being plated onto XLT-4 agar. All agar plates and broth tubes were incubated at 37 ^o^C for 24 – 48 h before colony forming units were enumerated. Reported counts were log-normalized and reported per gram PL dry weight. At least three single colonies representing three unique clones of *Salmonella* from each microcosm were randomly chosen from XLT-4 plate dilutions and archived in 30 % Luria Bertani (LB) glycerol at -80 ^o^C.

### DNA sequencing and bioinformatics

We sequenced 86 evolved *S*. Heidelberg isolates encompassing clones recovered from the four treatments. Isolates were selected randomly from the highest and lowest dilution of XLT-4 plates; and when populations decreased below limit of quantification (LOQ = 10 CFU/g dry weight of PL), isolates were selected after 24 h enrichment in BPW. Genomic DNA (gDNA) was extracted from pure isolate cultures using FastDNA Spin Kit (Mp Biomedicals, Solon, Ohio) and quantified with a Qubit Fluorometer (ThermoFisher Scientific). Whole genome sequencing (WGS) libraries were prepared using MiSeq Nextera XT library preparation kit (Illumina, Inc., San Diego, CA). Sequencing was performed on the Illumina MiSeq platform with 250-bp paired end reads using the MiSeq reagent V2 (500 cycles). All isolates were sequenced for an average coverage of 100X. Sequence reads were assembled de novo into contigs using SPAdes (23). The quality of assembled genomes was accessed using Quality Assessment Tool for Genome Assemblies (QUAST). (For detailed information on assembly, coverage and quality of sequenced genomes see Supplementary Files 1 and 2).

The original strains of SH-2813 and SH-116 are referred to as ancestral strains, while isolates from PL evolution experiments are referred to as evolved strains. Assembled contigs were submitted to the Center for Genomic Epidemiology’s multilocus sequence type (MLST) and PlasmidFinder modules to determine the whole-genome sequence types (ST) and existing plasmid replicon types. Prophages were identified using PHAST (24) and sequences for intact phages were extracted, and annotated with Prokka (25). The Comprehensive Antibiotic Resistance Database (CARD) was used for the identification of resistance genes present on plasmid and chromosome (26).

Plasmids were classified into MOB families using methods described previously (27). To investigate the topology of contigs identified with plasmid replicon, we used a custom script to inspect the path in the SPAdes FASTG assembly graph file that corresponded to the plasmid contig identified in the FASTA file; the contigs.paths file was used to determine correspondence between FASTA contigs and FASTG edges. Plasmid contigs were assigned as ‘complete circular’; ‘complete non-circular’, i.e. linear but not connected to other contigs; and ‘non-complete’, i.e. linear and connected to other contigs. (see Supplementary Methods for detailed information). Complete circular plasmids were annotated using Prokka (25) and mapped with Snapgene v 4.1.2. When coding DNA sequence (CDS) was annotated as hypothetical/uncharacterized, we performed a PSI-BLAST search with predicted protein sequences run to ≤3 iterations. Plasmids were analyzed by comparative analysis using ProgressiveMAUVE (28) and CDS predicted in regions with marked differences were aligned with MUSCLE and MAFFT (29, 30). Phylogenetic tree was produced from aligned conserved relaxase or mobilization protein sequences using the maximum likelihood method implemented in Randomized Axelerated Maximum Likelihood (RAxML) v 8 (31) using the PROTGAMMA/BLOSUM62 model and a rapid bootstrapping run to generate 1000 replicates. Samples with identical sequences were removed before phylogenetic tree was constructed. Trees generated were rooted with MbeA and MbeC sequences of *E. coli* (Accession number: NC001371).

### Single Nucleotide Polymorphism (SNPs) and InDel analysis

SNPs/InDels were determined by mapping raw sequence reads to the chromosome and plasmid genome of *S*. Heidelberg strain AMR588-04-00435 available in NCBI under accession numbers CP016573 and CP016575, respectively, using Burrows-Wheeler Aligner (BWA) (32). SAM file sorting, and removal of PCR duplicates was done using SAMtools (33). Genome Analysis ToolKit (34) with a minimum mapping quality of 30 and a minimum base quality of 30 was used for SNP identification. Manipulation of variant call format files (VCF) generated was done using vcftools v 0.1.12b (35). Shell script used and VCF files are available from the Dryad Digital Repository: https://doi.org/10.5061/dryad.pc6tp4q.

To obtain phylogenetic trees, SNPs and InDels mapping to plasmid sequences were removed and VCF files for SH-2813 and SH-116 evolved isolates were merged. The merged files were converted to a PHYLIP multiple alignment file using PGDSpider (36) and isolates with identical SNPs were removed before phylogenetic tree was constructed. The PHYLIP file was analyzed using maximum likelihood method implemented in RAxML v. 8 with the number of bootstrap replicates criteria set to extended majority rule consensus tree (`-N autoMRE`). We used the Jukes-Cantor (37) model of nucleotide substitution and GAMMA model of rate heterogeneity because they were the simplest model of sequence evolution predicted using jModelTest (38). Trees generated were rooted with either ancestral strain of SH-2813 or SH-116.

### Growth assay and plasmid copy number determination

Plasmid copy number (PCN) and differential expression of selected transcripts was determined by performing a growth assay on ancestral and evolved strains. Three populations were initiated from three single colonies from strains SH-2813_anc_, SH-2813_evol_, SH-116_anc_, and SH-116_evol_. An individual colony was transferred to 20 ml of Mueller Hinton Broth (MHB) in 50 ml centrifuge tube and incubated overnight at 37 ^o^C. Following overnight growth, 6 technical replicate populations (18 per strain) were established from each culture by transferring 3 ml to 50 ml centrifuge tubes containing 17 ml MHB. Cultures were incubated without shaking at 37 ^o^C for 24 h. Optical density (OD_600nm_) was periodically measured in the 3 populations with a Nanodrop 2000 Spectrophotometer (ThermoFisher Scientific). Bacteria concentration was determined for 2 populations at 0.5, 4 and 24 h via serial dilution plating on R2A agar. Additionally, bacteria culture (~ 20 ml) was filtered onto 0.2 μm pore size polycarbonate filters (MilliporeSigma, Burlington, MA) which were transferred to Lysing Matrix E and B tubes (MP Biomedicals, Solon, Ohio) for genomic DNA and RNA isolation respectively; and saved at 80 ^o^C until use.

Five primer sets specific to the glyceraldehyde-3-phosphate dehydrogenase, replication initiation protein, putative toxin-antitoxin, macrophage stimulating factor, a replication protein was designed and denoted *gapA,* repB_IncX1 Abi_ColE1-6kb, MSF_ColE1-4kb and rep respectively. The *gapA* is a single-copy gene of S. Heidelberg chromosomal DNA, while repB, Abi, MSF and rep are single-copy genes of IncX1, ColE1_6kb, ColE1_4kb and ColpVC_2kb, respectively. Real-time qPCR amplification and analysis were performed as previously described with a CFX96 Touch Real-Time PCR Detection System (Bio-Rad Inc., Hercules, CA). Specificity of primer pairs was verified by melting curve analysis and amplification factor for all assays was between 1.75 – 2.03 (Supplementary Table 9). PCN was determined as the copy ratio of plasmid encoded genes to *gapA* (39).

### Gene expression

For qRT-PCR, total RNA was isolated from membrane filters as described by ref (40) using FastRNA Pro Blue Kit (MP Biomedicals, Solon, Ohio). Purified RNA was treated with 4U DNA-free DNase (ThermoFisher Scientific, MA, USA) to remove genomic DNA contamination and quantified with a Qubit Flourometer (Thermo Fisher Scientific, MA, USA). DNase treated RNA was reversed transcribed with 100 U of high capacity cDNA reverse transcription kit (ThermoFisher Scientific, MA, USA) per manufacturer’s instructions.

Three primer sets targeting plasmid genes were designed in addition to the ones used for PCN determination. The primers target the toxin-antitoxin proteins (*relBE*) present on IncX1 and a putative aminoglycoside 6-N-acetyltransferase (aac-(6’)) present on ColE1_6kb. qRT-PCR was performed on generated cDNA as described for qPCR assay. Reaction mixtures (20 μl) for all assays contained 1X SsoAdvanced Universal SYBR Green Supermix (Bio-Rad Inc., Hercules, CA), 250 nM (each) primers, and 2 μl (20 -100 ng) of cDNA. Data were analyzed using the 2^-ΔΔCT^ method described by Livak and Schmittgen (41). For each plasmid target gene, the ratio of expression between ancestral and evolved strain was normalized to the expression of chromosome-encoded *gapA* and *gyrB* reference genes. The statistical significance of differences in gene expression relative to ancestor was determined using Wilcoxon signed-rank Test.

### Competitive fitness and conjugation assay

The fitness of three strains carrying the 3 Col plasmids (SH-2813_evol_, SH-116_anc_, and SH-116_evol_) was determined relative to SH-2813_nal_ nalidixic resistance with no Col plasmid. The SH-2813_nal_ strain is isogenic to SH-2813_anc_ but was made resistant to 200 ppm nalidixic acid by spontaneous mutation in *gyrA.* Five colonies from SBA for each strain were incubated in 9 ml of fresh MHB at 37 ^o^C for 20 h. Thereafter, cultures of the strains were mixed at a ratio of 1:1 (~500 μl of strain under study to 500 μl of SH-2813_nal_). We assumed that each strain had the same initial concentration because 24 h population was not significantly different (Table 3). Mixtures were diluted 10,000-fold in 50 ml centrifuge tubes (4 technical replicates per mixture) containing 20 ml of MHB and competed for 24 h at 37 ^o^C without shaking. The final population was measured by serial plating onto R2A agar supplemented with and without 32 ppm of nal. The fitness of the strains carrying Col plasmids relative to the SH-2813-nal strain was determined as described by ref (42):

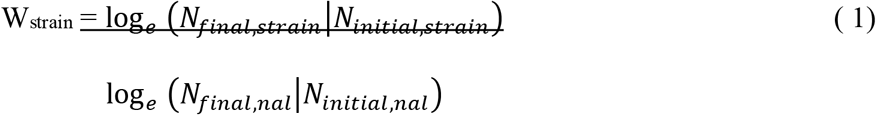

Where W_strain_ is the fitness of the strain carrying Col plasmids, *N_initial,strain_* and *N_final,strain_* are the numbers of cells (in colonies forming units (CFU)) of the strain carrying the Col plasmids before and after competition, and *N_initia,nal_* and *N_final,nal_* are the numbers of cells of SH-2813-nal before and after competition. To determine *N_final,strain_* we randomly screened 28 colonies from R2A without nal on R2A and Brilliant Green Sulfur (BGS) supplemented with 32 and 200 ppm of nalidixic acid, respectively. From here, the fractions of Col plasmids donor within each population was estimated by scoring the absence of growth on both R2A and BGS supplemented with nal. To assess possible fitness cost associated with *gyrA* mutation, SH-2813_nal_ was grown in single culture and plated on R2A with and without nal. The ratio of cells that grew on R2A without nal versus R2A with nal was 1.00 (s.e. =0.12, n =3).

To pool the information from the competition assay with conjugation assay we employed the same data set to determine the conjugation rate constant for the three Col plasmid types present during the competition assay. Conjugation rate constants were determined as described by Loftie-Eaton et al (43). Briefly, conjugation between plasmid-containing donors (D) and plasmid-free recipients (R) was measured under the same growth conditions used for the competition assay. To distinguish D and R, we randomly screened 56 colonies from R2A supplemented with 32 ppm nal for plasmid-specific gene. To do this, colonies were selected from the lowest and highest dilution plate (10^-5^ and 10^-7)^ for each strain of interest, and resuspended in 96-well PCR plates containing 100 μl DEPC-treated water. Thereafter a quick-boil DNA extraction at 95 ^o^C for 10 mins was done. Following the extraction, qPCR was used to estimate the proportion of positive “transconjugants” to Col plasmid-free recipients. The conjugation rate constant *γ* (in ml (c.f.u x *t*)^-1^) for the 3 Col plasmids in each competition assay was then determined using the following equation and as described by ref (43):

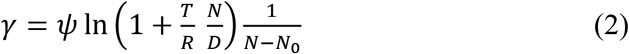

Where *ψ* is the maximum growth rate for the combined donor and recipient cultures, which is calculated from two bacteria concentration measurements taken during exponential phase and quantified on R2A without Nal, and calculated as follows:

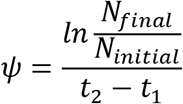

Total initial cell densities (*No*) and final (*N*) are the number of cells growing on R2A without nal at 0 and 24 h respectively. T is the number of transconjugants estimated from cells growing on R2A with nal.

### Antibiotic susceptibility testing

We performed antimicrobial susceptibility testing (AST) on selected evolved strains following National Antimicrobial Resistance Monitoring System (NARMS) protocols for Gram-Negative bacteria (44) and Kirby-Bauer disc diffusion assay (45). For more details on the disc diffusion protocol used, see Supplementary Methods.

For categorizing susceptibility, breakpoints were adopted from NARMs. There is no standardization by Clinical and Laboratory Standards Institute (46) for tobramycin, kanamycin, neomycin, netilmicin, and fosfomycin by disc diffusion for *S.* Heidelberg. Therefore, results were interpreted with diameter difference in zone of inhibition between isolates.

### Poultry litter DNA extraction

Two hundred and fifty milligrams of PL from each microcosm were transferred to PowerBead tubes in triplicate (Qiagen Inc, Germantown, MD) and stored at – 20°C until extraction, which occurred within 2 months of sampling. DNA was extracted using a Qiagen DNeasy Power-Soil DNA isolation kit, with the following modifications to the manufacturer’s instructions: (i) bead beating was conducted twice at 1800 rpm for 60 s, using a Fastprep-96 bead beater (MP Biomedicals, Solon, Ohio); and (ii) cleanup step with 800 μl of 100% ethanol was performed twice before cleanup with Solution C5. After elution of extracted DNA, an additional cleanup with Zymogen PCR inhibitor removal kit (Zymo Research Corp. Irvine, CA) was employed as per manufacturer’s instructions. Finally, all 3 purified DNA extracts from each microcosm was pooled and concentrated in an Eppendorf Vacufuge Plus Concentrator (Fisher Sci) at 30 ^o^C. Pelleted DNA was reconstituted in 100 μl 1X TE buffer (10 mM Tris-HCL, 1 mM EDTA, pH 7.5) and concentration was determined fluorometrically using Qubit Fluorometer (ThermoFisher Scientific, MA, USA). DNA was stored at -80 ^o^C until use.

### Statistical analysis

The two *S*. Heidelberg strains are the experimental units and PL represents the sampling unit for this study. Continuous variables did not meet the assumption of a normal distribution; therefore comparisons between strains or treatment were performed using Wilcoxon signed-rank Test. A principal component analysis (PCA) was performed on PL physico-chemical parameters (log-normalized) and *S*. Heidelberg concentrations to derive uncorrelated components representing most of the data information and analyze treatment effects on *S*. Heidelberg survival. A multiple linear regression was used to model the relationship between IncX1 and Col plasmid copy numbers derived from whole genome sequencing coverage. Analyses were performed using R (version 3.4.1).

### Ethics statement

The authors did not physically interact with chickens raised on the poultry litter used in this study, thereforewe were exempt from university guidelines and USDA/NIH regulations regarding animal use. Institutional Animal Care and Use Committee (IACUC) – UGA Office of Animal Care and Use.

### Data availability

All sequencing data are available under NCBI Sequence Read Archive (SRA) BioProject: PRJNA448609.

## RESULTS

### Experimental evolution system

The survival of two S. Heidelberg strains (SH-2813 and SH-116) in used PL was evaluated over a period of 14 days (Supplementary Fig. 1). We chose *S*. Heidelberg for its increasing role in food-borne outbreaks and its carriage of Col-like plasmids (14, 16, 17, 22, 47). The PL was previously used to raise 3 *Salmonella-free* flocks. In addition, 30 g of uninoculated PL was analyzed by culture method immediately after collection and at the start of the experiment (Day 0) for *Salmonella* isolation. No Salmonella was detected by direct culture on selective media or by enrichment of PL in BPW. The strains used were recovered from chicken carcass from two states in the USA and are closely related at the chromosomal level, differing from the *S.* Heidelberg reference genome used by only 109 and 89 SNPs/InDels. In addition, both strains carried an IncX1 plasmid and SH-116 carried 2 ColE1-like plasmids (4 and 6 kb) and a cryptic ColpVC plasmid (2 kb) (Table 1). The strains were chosen to investigate the role of Col-like plasmids on the fitness of S. Heidelberg in PL. We hypothesized that S. Heidelberg strains carrying Col plasmids will persist longer in PL.

The concentration of the two S. Heidelberg strains preconditioned in either PLE or BHIB decreased by several orders of magnitude in all microcosms and their survival dynamics differed by preconditioning media and strain-type (Fig. 1a).

**Figure 1.**
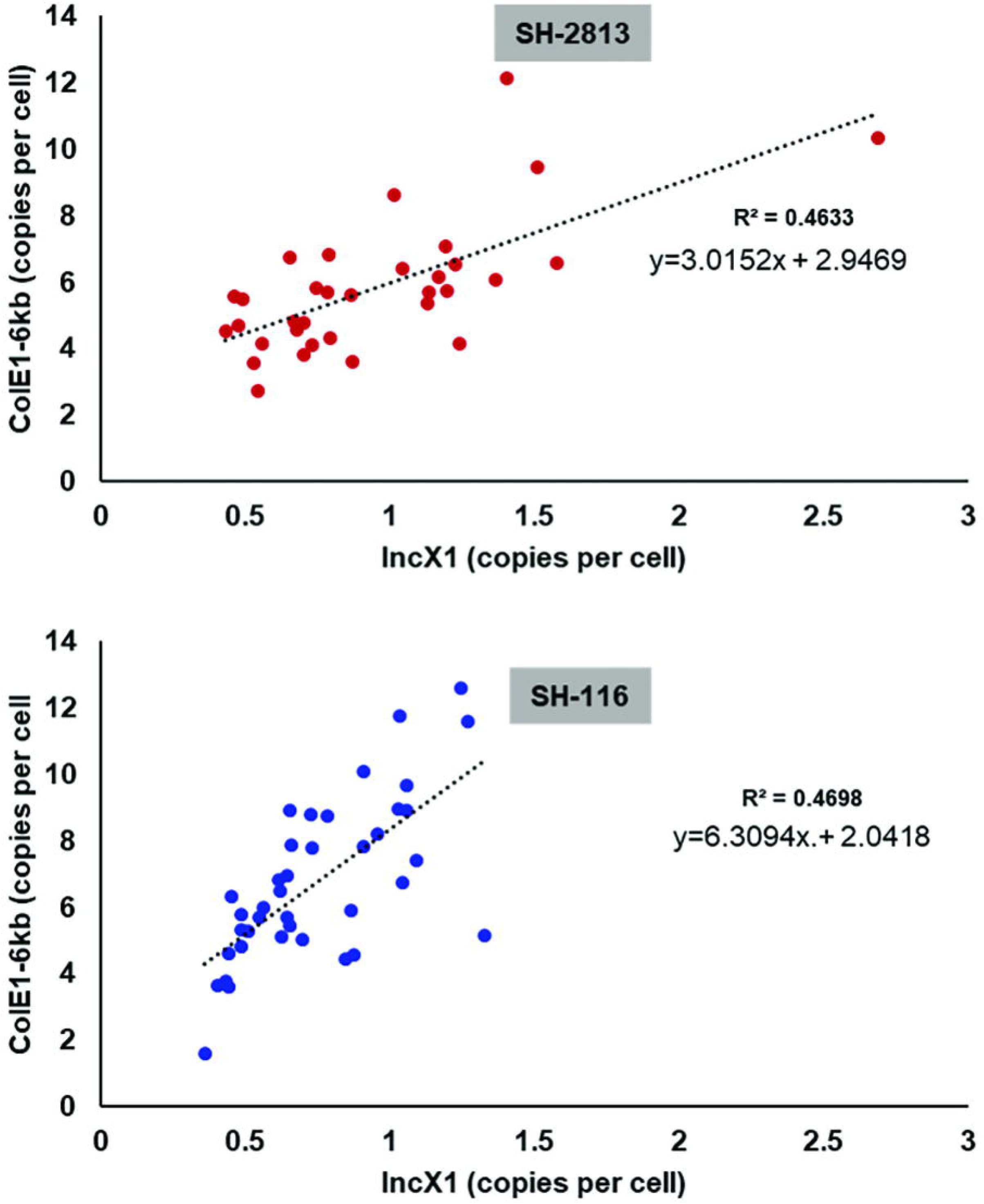
Survival curve and chromosomal mutation based phylogeny of evolved S. Heidelberg strains. (**a**) S. Heidelberg concentration from poultry litter microcosms reported in colony forming units (CFU) and normalized by litter dry weight. (**b**) Venn diagram showing the total number of chromosomal SNPs/InDels shared between isolates with hotspot mutations. (**c**) **Left.** Maximum-likelihood tree of SH-2813 and SH-116 isolates based on 213 SNPs/InDels. Clades and taxa highlighted in green denote isolates without hotspot mutations or Col plasmids, while clades highlighted with cyan boxes carry hotspot mutations and 3 Col plasmids. Taxa in parenthesis represent isolates with identical SNPs/InDels and not used in the reconstruction of the tree shown. Isolates with sample ID’s 1-24 are evolved isolates from strain SH-2813 and ID’s 25-48 are evolved isolates from strain SH-116. The numbers shown next to the branches represent the percentage of replicate trees where associated taxa cluster together based on ~200 bootstrap replicates. Numbers (1–18) denote arbitrarily assigned clade numbers. *Right*: Corresponding venn diagram comparing the total number of SNPs/InDels shared across time points. Tree was rooted with ancestral SH-2813.

The concentration of SH-2813-PLE, decreased by 4-log within 1 day of inoculation into PL but rebounded 1.5-log after 7 days. In contrast, the concentration of SH-2813-BHIB was below the LOQ within 1 day of inoculation into PL and increased approximately 4-log by day 7. The final concentration of SH-2813-PLE and SH-2813-BHIB in PL was log 2.5 ± 0.03 CFU g^-1^. On the other hand, the concentration of SH-116-PLE decreased gradually for 14 days while SH-116-BHIB was below LOQ within 1 day of inoculation into PL and increased by approximately 3-log at day 7 (Fig. 1a). The final concentration for SH-116 was below our LOQ for direct culture counts but the strain was isolated at varying levels following enrichment in buffered peptone water (BPW) (Supplementary Table 1).

We determined the pH, moisture, nutrients, and cations present in PL during the 14-day experiment (Supplementary Table 2). The litter pH and moisture level was 8.5 ± 0.23 and 32.3 ± 2 % at day 0 and 8.1 ± 0.09 and 16.7 ± 1 % at day 14. We observed significant differences for pH and moisture between PLE and BHIB microcosms (*p*. value <0.001; Wilcoxon signed-rank Test), with BHIB having higher moisture and pH (moisture =36 ± 2.1 %; pH= 8.75 ± 0.1) than PLE (moisture =25.4 ± 4 %; pH=8.17 ± 0.1) from day 0 to day 7. Also, percent reduction of total organic carbon (TOC), total carbon (TC), inorganic carbon (IC), total nitrogen (TN), ammonium (NH4), aluminum (Al), copper (Cu), Iron (Fe) and Zinc (Zn) in BHIB microcosms was higher than PLE microcosms after 14 days (Supplementary Table 2). To access the relationship between these physico-chemical parameters and *Salmonella* concentration, we performed a principal component analysis (PCA) on all variables measured. Two components of the PCA performed had eigenvalues higher than one and captured 68.8% of the variance in the data (Supplementary Fig. 2). Most variables were positively correlated with component 1 including pH (ρ = 0.79) and moisture (ρ = 0.87), with the exception of Mg, Ca and Mn that were negatively correlated with component 2.

We also determined the concentration of culturable total aerobic bacteria, coliforms and enterococci bacteria present in PL microcosms (Supplementary Table 1). Coliform bacteria were not recovered at any time point in our study, which agrees with studies showing low prevalence of Gram-negative bacteria in PL (49). SH-2813 microcosms had higher aerobic bacteria (log 7.6 ± 0.37 CFU g^-1^) than SH-116 (log 6.2 ± 0.07 CFU g^-1^).

### Whole genome sequencing

All isolates sequenced had an average coverage of 101 times with a range of 28 to 234 times Assembly size of sequenced genomes including extrachromosomal DNA ranged from 4.75 – 4.80 MB. Differences in genome size may be due to the presence or absence of mobile genetic elements including phages and plasmids. The average N50 for all isolates was 370, 097, which represents the average contig size after assembly with SPaDes assembler. Quality assessment and coverage report for all assembled genomes are available in Supplementary Files 1 and 2.

### Phylogenetic trees and chromosomal mutations

To investigate the genetics of *S*. Heidelberg persistence in PL, we isolated and performed WGS on 86 isolates obtained after 0, 1, 7 and 14 days in PL (Supplementary Table 3). First, we determined the genetic relatedness of the isolates recovered by reconstructing a maximum-likelihood tree (Fig.1c) with 213 SNPs/InDels identified in ancestral and evolved strains of SH-2813 and SH-116. We designated *S. Heidelberg* (CP013673) as the reference genome because it differed from the two strains by a low number of SNPs (<110) (Table 1). In addition, it carried similar IncX1 and ColpvC plasmids. After removal of identical SNPs/InDels, 28 isolates were not included in the reconstructed tree. All SH-2813 isolates recovered after 1 day of incubation in PL were represented by 4 clades highlighted in green (Fig. 1a). However, SH-2813 day 1 isolates recovered from PL that was previously enriched for 24 h in BPW (SH-2813BPW-Day1) were clustered with isolates recovered after day 7 and with all the SH-116 isolates (clades 5 -18). These isolates in clades 1-4 differed by 2 – 5 SNPs/ InDels from the ancestral SH-2813. A common mutation found between these isolates (9 of 12) was in the 23S-5S Internal Transcribed Spacer (ITS) region between positions 3503197 – 3503202 (Supplementary Table 4). Isolates not carrying this mutation (SH-5B, SH-13B, and SH-14B) had a mutation in the 16S rRNA region (position 4125232: A -> C). Mutation was also acquired in the 23S rRNA region (position 4129873; A -> G) for 6 isolates.

All SH-2813 isolates recovered after 7 days in PL (SH-2813-Day7) and SH-2813BPW-Day1 isolates (n =6) acquired a total of 76 new mutations that were absent in day 0 and 1 isolates. For SH-116, the number of SNP difference between two evolved isolates or the ancestral strain was not more than 3 SNPs at any time point. A total of 62 mutations were identified between SH-116, SH-2813-Day7 and SH-2813BPW-Day1 isolates (Fig. 1b and c). Of these mutations, 37 were SNPs/InDels present in the same loci on the chromosome and resulted in the same nucleotide change (Table 2).

**Table 2.** Mutations shared between evolved *S.* Heidelberg isolates.^a^.

| Loci on chromosome | Location <sup>b</sup> | Product | Mutation change <sup>c</sup> |
| --- | --- | --- | --- |
| 70249 | intragenic | Carbamoyl phosphate synthase small subunit | Asp--> Asp |
| 241459 | intragenic | Fimbrial protein (StfE) | Val--> Arg |
| 295492 | Intergenic | Upstream 16S ribosomal rRNA | N/A |
| 629229 | intragenic | Allantoin permease | Trp--> Stop |
| 774273 | intragenic | Amino acid ABC transporter permease | Ala --> Trp |
| 775920 | Intergenic | Upstream glutamate/aspartate ABC transporter | N/A |
| 823726 | Intergenic | Upstream peptide permease | N/A |
| 830662 | intragenic | UDP-galactopyranose mutase | Phe --> Leu |
| 832621 | intragenic | Galactosyl transferase (Pseudo) | ND |
| 832993 | intragenic | Galactosyl transferase (Pseudo) | ND |
| 1035518 | intragenic | Cysteine/glutathione ABC transporter permease (CydD) | Val --> Pro |
| 1103788 | Intergenic | Upstream FtsZ | N/A |
| 1210309 | intragenic | Proline dehydrogenase (PutA) | Arg --> Glu |
| 1288160 | intragenic | Ferric-rhodotorulic acid/ferric-coprogen receptor (FhuE) | Ala --> Arg |
| 1423283 | intragenic | Cyclic di-GMP regulator (CdgR) | Gln --> Stop |
| 1494796 | intragenic | Type III secretion system export apparatus protein (SsaV) | His --> Asn |
| 1628331 | intragenic | DUF1493 | ND |
| 1649579 | intragenic | NAD-dependent malic enzyme | Thr --> Thr |
| 1689446 | intragenic | 1-pyrroline dehydrogenase | Arg --> Thr |
| 2069216 | Intergenic | Upstream hypothetical protein | N/A |
| 2115173 | Intergenic | Upstream cobyrinic acid a,c-diamide synthase | N/A |
| 2333347 | intragenic | Acetylase | ND |
| 2517571 | Intergenic | Upstream outer membrane protease (OmpT) | N/A |
| 2757879 | IntraRNA | 5S ribosomal rRNA | N/A |
| 2880324 | intragenic | Ribonucleotide-diphosphate reductase subunit alpha (NrdA) | Asp --> Ala |
| 3212823 | intragenic | tRNA (guanosine(46)-N7)-methyltransferase (TrmB) | Arg --> Ala |
| 3324819 | intragenic | Transcriptional regulator (PadR) | Leu --> Gln |
| 3336383 | intergenic | Upstream hypothetical protein | N/A |
| 3357216 | intragenic | Transcriptional regulator (TdcA) | Phe --> Glu |
| 3523467 | Intragenic | 50S ribosomal protein L15 | Ala --> Arg |
| 3594367 | intragenic | Two-component system sensor histidine **kinase (EnvZ)** | Val --> Asn |
| **3629811** | **intragenic** | **2-ketogluconate transporter** | **Ser --> Ser** |
| **3791102** | **intragenic** | **3-dehydro-L-gulonate 2-dehydrogenase** | **Gly --> Ser** |
| **4265903** | **intragenic** | **TonB-dependent vitamin B12 receptor (BtuB)** | **Ala --> Ser** |
| **4303417** | **intragenic** | **Hypothetical protein** | **Asp --> Ala** |
| **4607527** | **intragenic** | **Transposase (IS200)** | **Glu --> Val** |
| **4652337** | **intergenic** | **Upstream hypothetical protein** | **N/A** |
| 295539 | Intergenic | Upstream rRNA-16S ribosomal RNA | N/A |
| 295548 | Intergenic | Upstream rRNA-16S ribosomal RNA | N/A |
| 295551 | Intergenic | Upstream rRNA-16S ribosomal RNA | N/A |
| 295555 | Intergenic | Upstream rRNA-16S ribosomal RNA | N/A |
| 295561 | Intergenic | Upstream rRNA-16S ribosomal RNA | N/A |
| 295564 | Intergenic | Upstream rRNA-16S ribosomal RNA | N/A |
| 926872 | Intragenic | hypothetical protein<sup>#</sup> | Leu --> Asp |
| 1367477 | Intragenic | aldo/keto reductase<sup>^</sup> | Thre --> Ile |
| 1420609 | Intragenic | vitamin B12 ABC transporter permease (BtuC) | Val --> Gly |
| 2018078 | Intergenic | Upstream methyl-accepting chemotaxis protein II<sup>#</sup> | N/A |
| 2049907 | Intragenic | flagellin (FliC)<sup>^</sup> | Leu --> Ala |
| 2115448 | Intragenic | hypothetical protein<sup>#</sup> | Leu --> Tyr |
| 2163440 | Intragenic | Phosphomannomutase<sup>#</sup> | Gly --> Thr |
| 2307329 | Intragenic | bifunctional PTS fructose transporter subunit IIA/HPr<sup>#</sup> | Ala --> Val |
| 2978093 | Intragenic | DNA mismatch repair protein (MutS)<sup>#</sup> | Met--> Asn |
| 3503198 | ITS | 23S -5S rRNA ITS<sup>^</sup> | N/A |
| 3503200 | ITS | 23S -5S rRNA ITS<sup>#</sup> | N/A |
| 3503201 | ITS | 23S -5S rRNA ITS<sup>*</sup> | N/A |
| 3503201 | ITS | 23S -5S rRNA ITS<sup>*</sup> | N/A |
| 3503201 | ITS | 23S -5S rRNA ITS<sup>*</sup> | N/A |
| 3503213 | ITS | 23S -5S rRNA ITS<sup>^</sup> | N/A |
| 3503219 | ITS | 23S -5S rRNA ITS<sup>^</sup> | N/A |
| 3532866 | Intergenic | Upstream 30S ribosomal protein S10 and leader peptidase hopD<sup>#</sup> | N/A |
| 3947635 | Intragenic | Cytochrome c biogenesis protein (CcmH)<sup>^</sup> | ND |
| 4035545 | ITS | 23S -5S rRNA ITS | N/A |
| 4035546 | ITS | 23S -5S rRNA ITS | N/A |
| 4035547 | ITS | 23S -5S rRNA ITS<sup>^#</sup> | N/A |
| 4035547 | ITS | 23S -5S rRNA ITS<sup>^#</sup> | N/A |
| 4035548 | ITS | 23S -5S rRNA ITS<sup>^</sup> | N/A |
| 4035549 | ITS | 23S -5S rRNA ITS | N/A |
| 4035551 | ITS | 23S -5S rRNA ITS | N/A |
| 4035556 | ITS | 23S -5S rRNA ITS | N/A |
| 4035560 | ITS | 23S -5S rRNA ITS* | N/A |
| 4035560 | ITS | 23S -5S rRNA ITS* | N/A |
| 4035561 | ITS | 23S -5S rRNA ITS | N/A |
| 4035562 | ITS | 23S -5S rRNA ITS* | N/A |
| 4035562 | ITS | 23S -5S rRNA ITS* | N/A |
| 4035566 | ITS | 23S -5S rRNA ITS | N/A |
| 4035567 | ITS | 23S -5S rRNA ITS | N/A |
| 4035570 | ITS | 23S -5S rRNA ITS<sup>^</sup> | N/A |
| 4035573 | ITS | 23S -5S rRNA ITS | N/A |
| 4035575 | ITS | 23S -5S rRNA ITS<sup>^</sup> | N/A |
| 4035576 | ITS | 23S -5S rRNA ITS* | N/A |
| 4035576 | ITS | 23S -5S rRNA ITS* | N/A |
| 4271745 | IntraRNA | rRNA-23S ribosomal RNA<sup>#</sup> | N/A |
| 4271746 | IntraRNA | rRNA-23S ribosomal RNA<sup>#</sup> | N/A |
| 4271748 | IntraRNA | rRNA-23S ribosomal RNA<sup>#</sup> | N/A |
| 4271751 | IntraRNA | rRNA-23S ribosomal RNA<sup>#</sup> | N/A |
| 4289060 | Intragenic | DNA-directed RNA polymerase subunit beta (RpoC) <sup>#</sup> | Ile --> Asn |
| 4366478 | Intragenic | Maltose ABC transporter substrate-binding protein (MalE) <sup>#</sup> | Gly --> Ser |
| 4482602 | Intergenic | Upstream acid phosphatase<sup>^</sup> | N/A |

Note.
<sup>a</sup> Mutations are present in Col-plasmid carrying isolates and absent in isolates without Col plasmids. SNPs/InDels were compared to reference genome CP016573.
<sup>b</sup> Intragenic- present within a coding sequence, Intergenic – non coding region between two genes, ITS – internal transcribed spacer, IntraRNA – present within non-coding RNA.
<sup>c</sup> N/A – not available (SNP/InDel present in non-coding sequence), ND- not determined.
**Boldness** denotes hotspot mutations present in all Col-plasmid carrying isolates (n=37).
<sup>#</sup> Single nucleotide variants present in <1 % of SH-2813 isolates (n=14).
<sup>^</sup> Single nucleotide variants present in <1 % of SH-116 isolates (n=12).
\* “Heterozygous” SNPs/InDels present in multicopy gene.

These mutations were in proteins or intergenic regions associated with diverse processes such as carbomoyl phosphate synthesis, motility (StfE), galactosyltransferease (a pseudogene), two-component system sensor histidine kinase (EnvZ), cyclic -di-GMP synthetase (c-di-GMP), SPI-2 type III secretion system apparatus protein (SsaV), siderophore transport (BtuB and FhuE), cysteine/glutathione transporter (CydD), 50S ribosomal protein (L15) and a protein of unknown function (DUF1493) etc.

Twenty-five of the 62 mutations were present in ~ 70% of SH-2813 and SH-116 isolates (Table 2). The majority of these mutations were multiple SNPs/InDels present in the 23S-5S ITS region between positions 4033545 – 4035576 on the chromosome. Further, 14 single nucleotide variants (SNVs) were present only in a handful of SH-2813 isolates and absent in all SH-116 isolates. For instance, strain SH-2813-17BBPW carried 5 unique SNVs including a non-synonymous mutation (met1asn) in the DNA mismatch protein (MutS) making it a spontaneous mutant (Table 2; Supplementary Table 4). For SH-116, 11 unique SNVs were identified in 23S-5S ITS region (n =8) and in proteins encoding aldo/keto reductase, phase 1 flagellin (FliC) and cytochrome c biogenesis protein (CcmH).

### IncX1 and Col plasmids are associated with *S*. Heidelberg persistence in PL

We next determined if S. Heidelberg isolates recovered from PL acquired new plasmids. All SH-2813_evol_ isolates (n= 27) obtained from day 7 and 14 microcosms and SH-2813BPW-Day1 isolates carried 3 Col plasmids in addition to an IncX1 plasmid. For SH-116, IncX1 and Col plasmids were present in all sequenced strains (n = 38) regardless of time of incubation in PL. The majority (276/306) of the plasmid contigs represented complete circular plasmids (Supplementary Table 5). The IncX1 belonged to the MOB_Q_ family of relaxases and was present in all sequenced genomes. The ColE1 plasmids belonged to the MOBp family and showed a conserved size of 6189 (ColE1-6kb) and 4784 (ColE1-4kb) bp respectively. The cryptic ColpVC was 2223 bp (ColpVC-2kb). For all complete IncX1 plasmids, there was a 21-base pair (bp) difference in strains carrying the Col plasmids (37, 824 bp) compared to strains with no Col plasmids (37, 845 bp).

### Direct-repeat deletion is associated with IncX1 copy number increase

To determine the effect of the 22-bp deletion identified in the IncX1 plasmids present in S. Heidelberg populations carrying Col plasmids, all complete circular IncX1 plasmid sequences were subjected to whole genome sequence alignment and SNV identification. The 22-bp direct-repeat deletion was located at the origin of replication (ori) adjacent to the replication protein (RepB) and an additional 1-bp (G-> GA) insertion occurred in the type IV secretion protein (T4SS) VirB8 (Fig. 2a and Supplementary Table 5).

**Figure 2.**
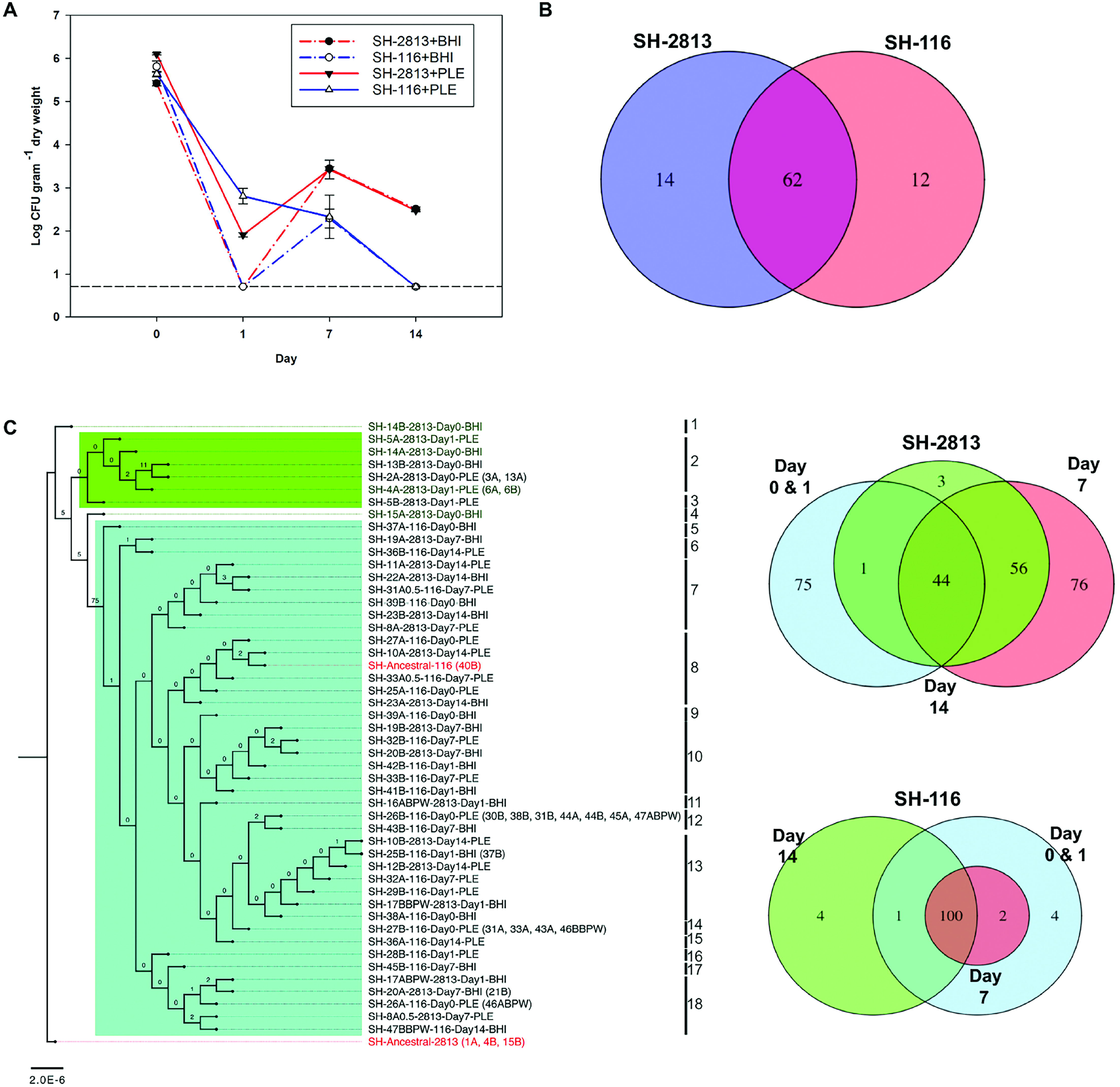
Deletion of 22-bp iteron of IncX1 is associated with a reduction in *rep* expression. (**a**) Annotated IncX1 plasmid map of SH-2813_anc_ and SH-116_anc_. Regions that acquired mutations in evolved strains carrying Col plasmids are highlighted in rectangular dashed boxes (**b**) Multiple alignment of the IncX1 iteron-based direct repeat (DR) origin of selected strains from this study, and 22 S. Heidelberg genomes available on NCBI. (**c**) Differences in expression of IncX1 genes in evolved SH-2813 and SH-116 relative to SH-2813_anc_ and SH-116_anc_ (log2 scale, N =3, \**p*.value =0.086 (Wilcoxon signed-rank Test). The error bars represent the standard error of the mean.

The 22-bp deletion reduced the number of direct repeats from 3 to 2, making it a truncated *ori* (Fig. 2b). Direct repeat iterons of theta plasmids are known to bind Rep proteins and have been shown to participate in negative control of their initiation (50). Deleting or adding more copies of the iteron have been shown to increase or decrease plasmid copy number (PCN) (51). Therefore, we hypothesized that the direct repeat deletion in IncX1 of SH-2813_evol_ would result in higher copies compared to SH-2813_anc_ since the 22-bp is present in the ancestral strain while no change would be expected in IncX1 copies for SH-116_evol_. To test this hypothesis, we determined the copy numbers of IncX1 and Col plasmids present in ancestor (SH-2813_anc_ and SH-116_anc_) and 2 evolved strains (SH-2813_evol_ and SH-116_evol_ respectively). We assumed that these evolved strains represent the “fittest” SH-2813 and SH-116 strains given their persistence in PL microcosms for 14 days.

As expected, the copy number of IncX1 for SH-2813_evol_ (~7 copies/cell; s.d.=1.8) was higher than that of SH-2813_anc_ (~4 copies/cell; s.d.=0.3) at 0.5 h (*p*. value = 0.086; Wilcoxon rank-signed test). In contrast, SH-116 did not exhibit any significant difference in IncX1 copies between SH-116_anc_ (~5 copies/cell; s.d.=1.7) and SH-116_evol_ (~6 copies/cell; s.d.=0.8) (*p.* value = 0. 773) (Table 3).

**Table 3.** Plasmid copy numbers of ancestral and evolved strains.

|  |  |  |  | Plasmid Copy Number/Cell (S.D) |  |  |  |
| --- | --- | --- | --- | --- | --- | --- | --- |
| Strain | Time (h) | Optical Density (S.D) | Bacteria concentration (Log CFU/ml) | IncX1 | ColE1-6kb | ColE1-4kb | ColpVC |
| SH-2813<sub>anc</sub> | 0.5 | 0.015±0.00 | 7.98±0.0 | 4.17±0.3 | N/A | N/A | N/A |
|  | 4 | 0.040±0.00 | 8.60±0.2 | 0.47±0.5 | N/A | N/A | N/A |
|  | 24 | 0.070±0.00 | 8.90±0.1 | 0.97±0.9 | N/A | N/A | N/A |
| SH-2813<sub>evol</sub> | 0.5 | 0.016±0.00 | 7.94±0.0 | 6.92±1.8 | 10.71±4 | 12.4±5.5 | 110.77±45 |
|  | 4 | 0.056±0.00 | 11.16±2.1 | 0.59±0.3 | 3.28±0.6 | 2.77±1.0 | 86.21±25 |
|  | 24 | 0.080±0.00 | 8.57±0.0 | 0.64±0.1 | 5.25±0.7 | 4.2±0.6 | 115.32±70 |
| SH-116<sub>anc</sub> | 0.5 | 0.014±0.00 | 7.83±0.1 | 4.55±1.7 | 0.27±0.3 | 2.54±1.9 | 82.39±8 |
|  | 4 | 0.037±0.00 | 8.56±0.1 | 0.80±1.3 | 0.14±0.0 | 0.54±0.7 | 54.01±26 |
|  | 24 | 0.093±0.00 | 9.07±0.0 | 2.33±9.3 | 2.23±0.0 | 0.92±1.3 | 39.94±28 |
| SH-116<sub>evol</sub> | 0.5 | 0.017±0.00 | 7.91±0.0 | 5.61±0.8 | 9.44±3.5 | 10±4.4 | 90.47±45 |
|  | 4 | 0.065±0.00 | 8.73±0.2 | 0.60±0.8 | 3.83±1.7 | 2.9±1.3 | 93.83±11 |
|  | 24 | 0.069±0.00 | 8.69±0.1 | 0.35±0.4 | 4.39±1.0 | 3.5±0.3 | 72.56±20 |

Note. –
SH-2813<sub>anc</sub> - Original SH-2813 strain before inoculation into PL, SH-2813<sub>evol</sub> - SH-2813-12A-day14-PLE, SH-
116<sub>anc</sub> – Original SH-116 strain before inoculation into PL, SH-116<sub>evol</sub> = SH-116-36B0.5-day14-PLE.
N/A= not applicable

### Gene expression of IncX1 replication protein and stabilization genes

We monitored the expression of the *repB* replication gene and *relBE* genes which encode a type II toxin-antitoxin (TA) system associated with the IncX1 plasmid. The *relBE* genes were selected since they are adjacent to the *ori* (Fig. 2a) suggesting they may share the same promoter as *repB*; and due to their implicated role in plasmid stabilization (52, 53). Gene expression results in SH-2813_evol_ revealed a marked 4.5-fold reduction in *repB* expression and 2.2-fold reduction in *relB* expression at 4 h relative to SH-2813anc; while the expression levels of toxin *relE* remained unchanged in both ancestral and evolved SH-2813 (Fig. 2c).

SH-116 strains lacks the extra 22-bp direct repeat present on IncX1, thus we expected no change in Rep expression. However, there was a change in expression for these transcripts for SH-116_evol_ — reductions in *repB* and *relB* and no change in *relE* when compared to SH-116_anc_ (Fig. 2c). We speculate that the reduction in expression of Rep for SH-116 evolved strains was a result of carrying higher copy ColE1 plasmids than SH-116_anc_. In fact, SH-116_evol_ carried ColE1 plasmids with higher copies (ColE1-6kb =9±3.5, ColE1-4kb = 10±4.4, ColpVC-2kb = 90±45) than SH-116_anc_ (ColE1-6kb =0.3±0.3 ColE1-4kb = 3±1.9, ColpVC-2kb = 82±7.9) (Table 2). The presence of this high copy ColE1 plasmids correlated with faster growth rates in MHB. The optical density of SH-2813_evol_ and SH-116_evol_ was 1.4 and 1.7-fold higher than that of SH-2813_anc_ and SH-116_anc_, respectively, at 4 h (Table 3). This difference in copy number was further corroborated by comparing PCN determined from WGS assembly coverage, showing that all SH-116_evol_ isolates from day 1, 7 and 14 on average had higher ColE1 plasmid copies than SH-116_anc_ (Supplementary Table 6). Finally, a multiple regression analysis on IncX1 and Col PCN showed that IncX1 was a statistically significant predictor of Col plasmid copies in a cell (F (3, 68) =13.82, *p*.value < 0.000, R^2^ =0.39). SH-116 strains were predicted to have approximately six ColE1-6kb plasmid copies for every IncX1, whereas, SH-2813 was predicted to have 3 copies per IncX1 (Fig. 3).

**Figure 3.**
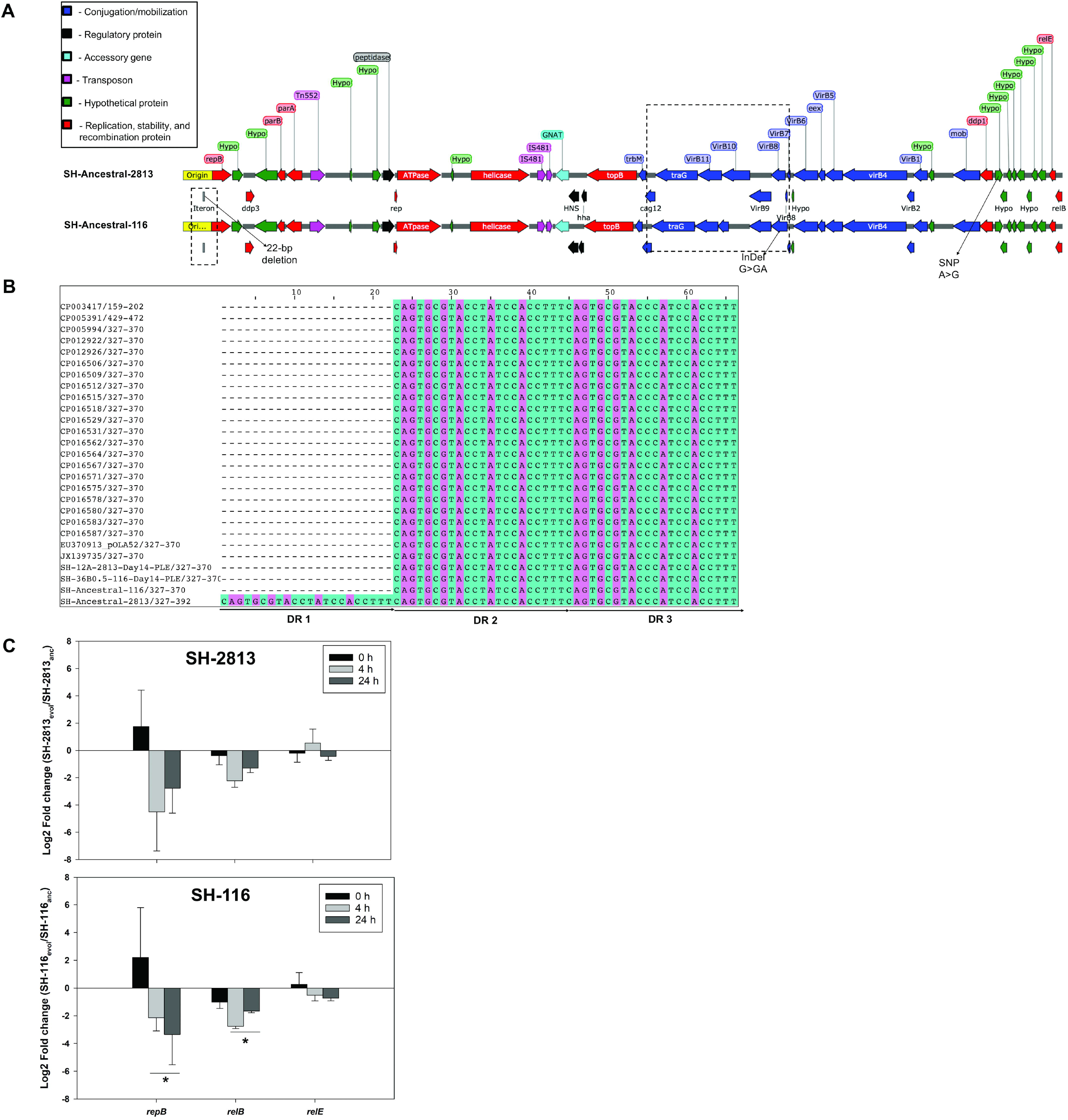
Correlation between IncX1 and ColE1-6kb plasmid copies. Plasmid copy number (PCN) was determined from whole genome sequencing de-novo assembly coverage (see Table S6.

### ColE1 plasmids differed in mobilization genes and non-coding RNA

To determine if there are genetic differences between Col plasmids carried by SH-116 and SH-2813, we used ProgressiveMauve (28) to perform multiple whole genome alignments on DNA sequences from complete Col plasmids. Results showed that ColE1 plasmids present in sequenced strains were highly similar and only differed in nucleotide regions coding for mob genes (Fig. 4; Supplementary Fig. 3b and c).

**Figure 4.**
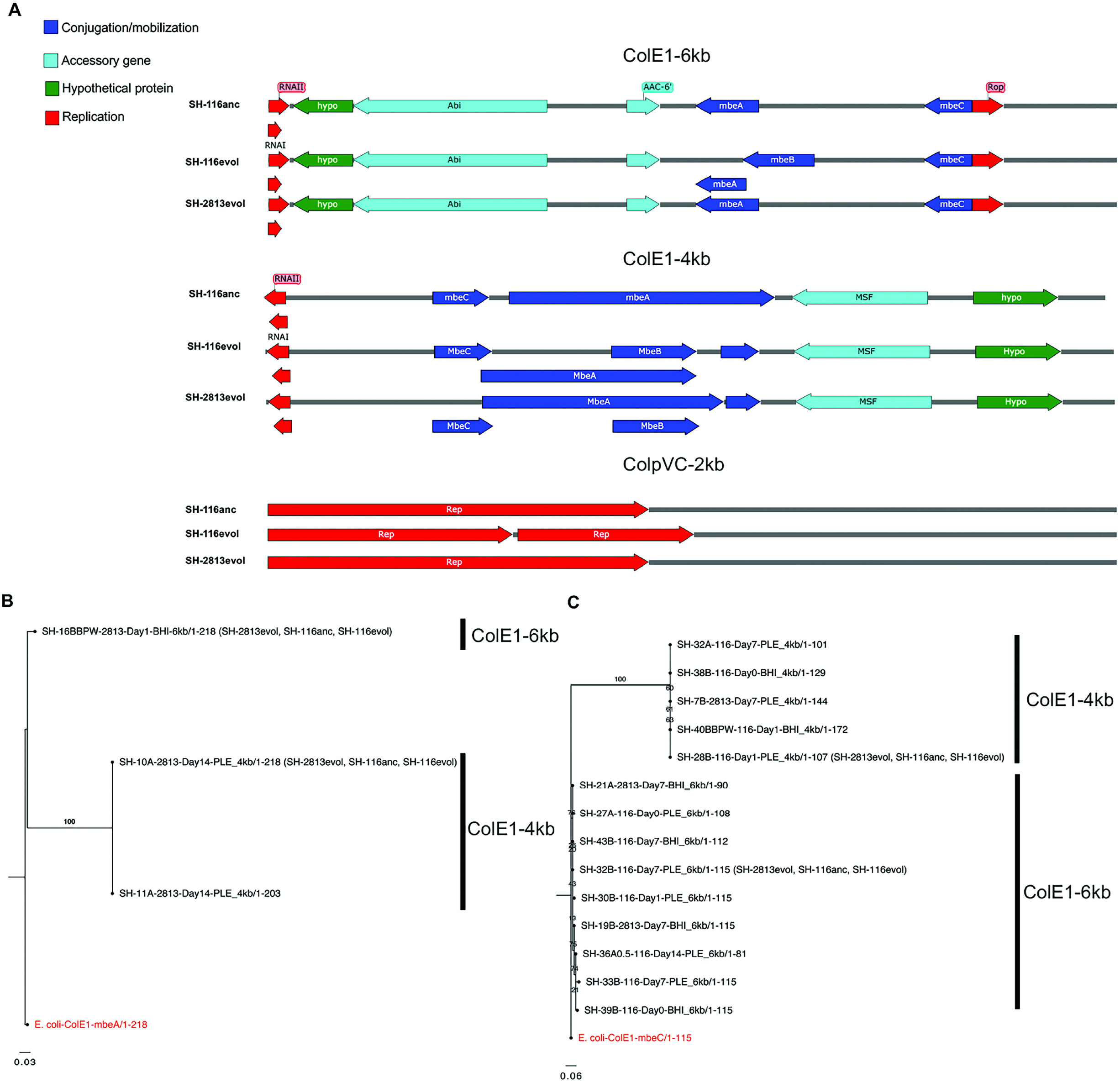
Linear maps of Col plasmids and phylogeny based on conserved mobilization genes. (**a**) Annotated maps of Col plasmids carried by ancestral and evolved isolates of *S*. Heidelberg used for competition experiments. (**b and c**) Maximum-likelihood based tree of aligned conserved protein sequences of (**b**) relaxase (MbeA) and (**c**) MbeC genes of Col plasmid-carrying isolates. Plasmids with identical MbeA (n=106) and MbeC (n=112) sequences were not included in the reconstructed tree. The numbers shown next to the branches represent the percentage of replicate trees where associated taxa cluster together based on 1000 bootstrap replicates. Taxa in parenthesis represent selected isolates used for competition experiments. Trees were rooted with MbeA and MbeC sequences of *E. coli* (NCBI accession number: NC_001371).

We identified 5-6 open reading frames (ORF) encoding 2-3 mobilization (mob) genes, a putative type IV TA system (Abi), a putative aminoglycoside 6-N-acetyltransferase (aac(6’)-Ib) and a hypothetical protein for ColE1-6kb (Fig. 4). The ColE1-4kb plasmids had 4-6 ORFs encoding a protein named macrophage stimulating factor (MSF), a hypothetical protein and 1-4 mob genes. Both ColE1 plasmids harbored a non-coding RNAI and RNAII, while ColE1-6kb carried an additional non-coding repressor of primer (Rop) protein known to regulate replication and ColE1 plasmid copy numbers. The ColpVC-2kb plasmids had 2-3 ORFs, encoding 1-2 replication proteins (Rep) and a hypothetical protein (Fig. 4). The RNAI present in the ColE1 plasmids had approximately 62 % DNA sequence homology (data not shown).

These mob CDS were identified as MbeA and MbeC and were present in different sizes and arrangement. In some cases, the *mbeA* gene overlapped ORF annotated as *mbeB and mbeD* (Fig. 4), suggesting MbeB and MbeD was part of the mob region in some but not all ColE1 plasmids. To determine if there are differences in these proteins between Col plasmids, we performed a multiple alignment of conserved MbeA relaxase and MbeC proteins. We observed high conservation in the N-terminal region and high variation in their C-terminal (data not shown). According to the phylogenetic tree reconstructed for these conserved amino acids, MbeA proteins belonging to ColE1-4kb plasmids clustered in a unique clade, whereas the ColE1-6kb was placed closer to the signature *E. coli* ColE1 plasmid (NC_001371) (Fig. 4b). After the removal of identical sequences, only MbeA sequences belonging to 3 SH-2813 isolates were included in the final tree The MbeA tree constructed for ColE1-6kb and ColE1-4kb plasmids were represented by 1 and 2 isolates, respectively. On the other hand, the two clades observed for MbeC tree had 9 and 5 isolates for ColE1-6kb and ColE1-4kb plasmids (Fig. 4c).

### Col plasmids did not impose a fitness cost on *S*. Heidelberg

Although acquisition of Col plasmids could result in faster growth in host bacterium it may also translate to a fitness cost if their regulation is not equally constrained. To establish if acquisition of Col plasmids imposed a fitness cost on S. Heidelberg, we competed 2 evolved (SH-2813_evol_ and SH-116_evol_) and 1 ancestral (SH-116_anc_) strain carrying three Col plasmids against a Col plasmid-free isogenic SH-2813_anc_ harboring a *gyrA* mutation for nalidixic acid resistance (SH-2813_nal_). The fitness of each strain was determined relative to SH-2813_nal_ as described by Millan et al (42). Results show that Col plasmid carrying strains had a higher relative fitness of 1.27 (s.d. =0.12, n =3) compared to SH-2813_nal_, and differences were observed between strains (Fig. 5a).

**Figure 5.**
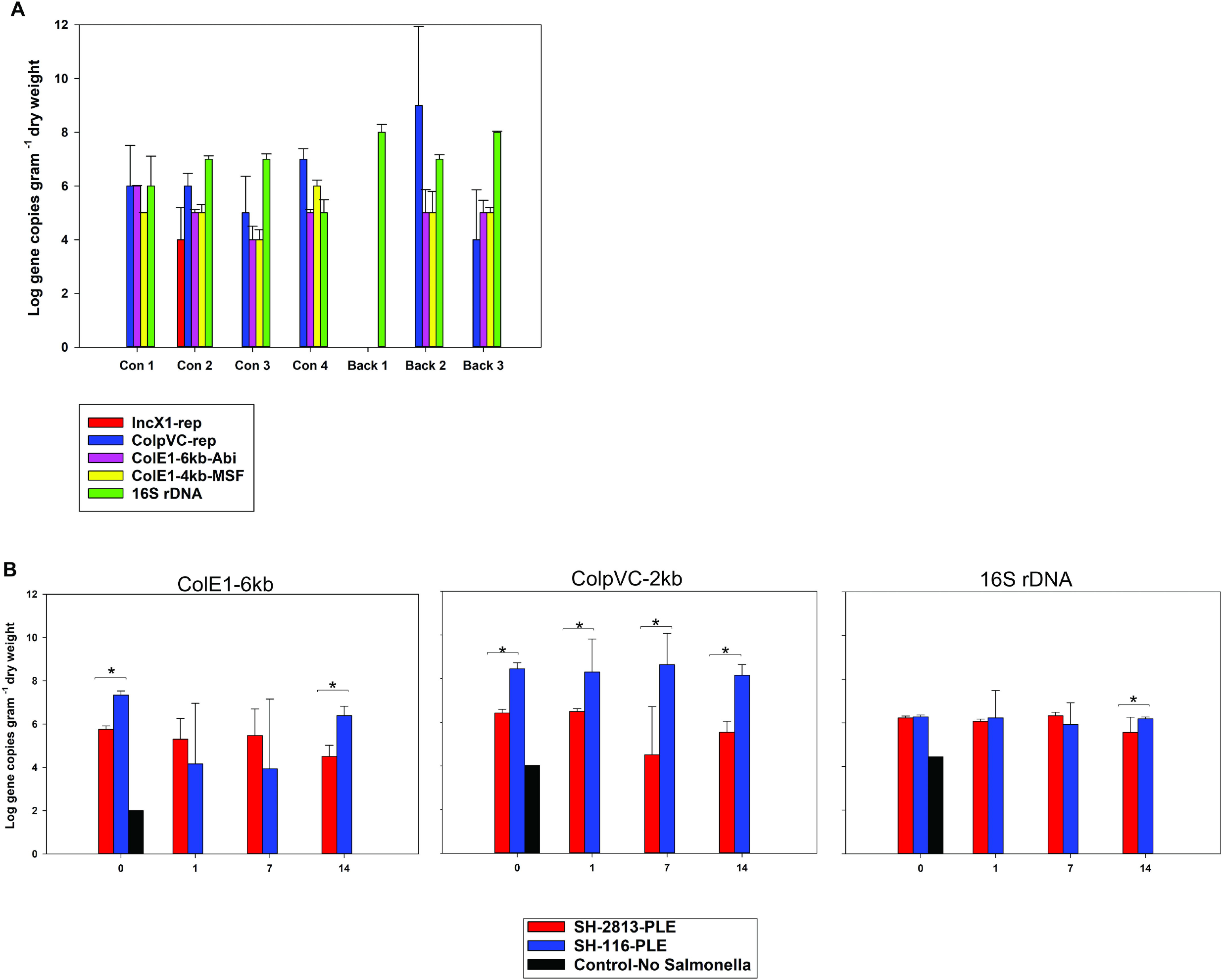
Fitness and Conjugation rates of evolved S. Heidelberg isolates. **(a)** Fitness of Col plasmid-carrying strains of SH-2813 and SH-116 relative to a Col plasmid-free ancestor of SH-2813_nal_. Bars represent means of four technical replicates for each population. The horizontal dashed red line represents the fitness of SH-2813_nal_. **(b)** Conjugation rates of Col plasmids from Col plasmid-carrying strains to SH-2813_nal_ ancestor. Conjugation rates were estimated using data from fitness experiment. Bars represent means of four technical replicates for each population. The transfer rate for ColE1-4kb plasmid from SH-116_evol_ to SH-2813_nal_ was not determined. Symbols: Abortive phage infection protein of ColE1-6kb (Abi-ColE1-6kb), macrophage stimulating factor protein of ColE1-4kb (MSF-ColE1-4kb) and replication protein of ColpVC (rep-ColpVC).

The transfer rate for ColE1-4kb plasmid from SH-116_evol_ to SH-2813_nal_ was not determined because none of the transconjugants screened by qPCR was positive for the MSF gene harbored on this plasmid. Consequently, the differences in fitness between strains may be attributed to a higher conjugation rate observed for ColE1-6kb and ColpVC for SH-116_evol_ (Fig. 5b).

### Antimicrobial resistance associated with the acquisition of Col plasmids

The ColE1-6kb plasmid in the current study carries a putative aac (6’)-Ib gene that is 56 % (*e.value* = 1<e-13) identical to the aac (6’)-Ib gene present in *Klebsiella pneumoniae* Sul1-type integron (GenBank accession number: AAR18814.1) and 34 % (*e.value* = 6<e-28) identical to the one present in *Thiomona* sp. X19 (GenBank accession number SCC93141.1); suggesting it may play a role in aminoglycoside resistance (Fig. 6a and b).

**Figure 6.**
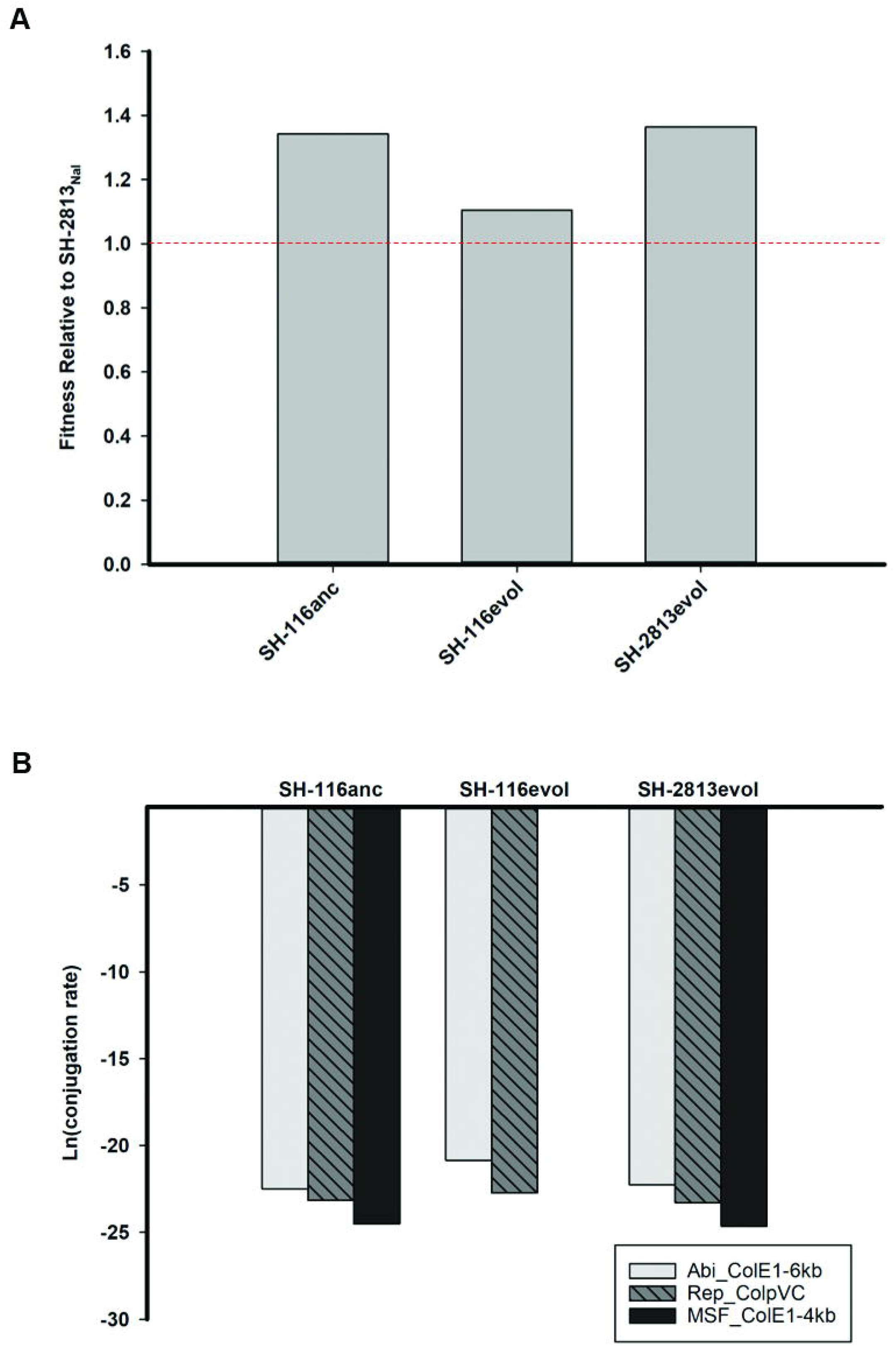
Acquisition of ColE1 plasmids was associated with decreased susceptibility to aminoglycosides and fosfomycin. (a) Multiple alignment of conserved N-terminal residues for aminoglycoside acetyltransferase (AAC-6’) genes encoded in the *S*. Heidelberg (SH-2813) chromosome and ColE1-6kb plasmid from this study, and AAC-6’ genes with known resistance phenotype. (b) Corresponding neighbor-joining tree of aligned AAC-6’ protein sequences. (c) Inhibition zone diameter (mm) distributions for evolved strains and selected antimicrobial drugs. SH-2813 (n= 27) and SH-116 (n= 18) isolates tested were recovered after 0 and 14 days of microevolution in PL. A decrease in inhibition zone corresponds to a decrease in susceptibility to selected antibiotic.

Further, *S*. Heidelberg ancestral strains each carried chromosomally encoded aminoglycoside (aac (6’)-Iy) and fosfomycin (fosA7) genes (54) that have been demonstrated to confer resistance to tobramycin, netilmicin, neomycin, amikacin, kanamycin and fosfomycin (55–57). To assess the correlation between AMR and the acquisition of Col plasmids, we performed antibiotic susceptibility testing (AST) on evolved S. Heidelberg strains (n=42) on the NARMs antimicrobials (44). In addition, we used disc-diffusion assay (45) to evaluate susceptibility to tobramycin, netilmicin, gentamicin, neomycin, kanamycin and fosfomycin.

The majority of the isolates tested (97.6 %) were susceptible to the 14 drugs on NARMs AST panel (Supplementary Table 7). However, 12 of 16 (75 %) SH-2813_evol_ isolates recovered from PL on day 7 and day 14 displayed an increase in MIC for gentamicin. The average zone of inhibition (mm) for kanamycin, tobramycin and fosfomycin for SH-2813_anc_ strains was larger than the zones measured for SH-2813_evol_ strains (Wilcoxon signed-rank test; p-value < 0.0001) (Fig. 6c). In contrast, SH-116 showed no significant difference in zone of inhibition between SH-116_anc_ and SH-116_evol_ strains for any of the antibiotics tested (Wilcoxon signed-rank test; p-value > 0.01).

### Col plasmid-encoded genes are prevalent in PL

To extrapolate beyond our microcosm evolution experiment, bioinformatics and quantitative PCR were used to assess the presence of these Col plasmids across sequenced S. Heidelberg genomes and in various types of PL. DNA sequences for complete Col plasmids from this study were subjected to BLAST searches against the non-reduntant (nr) GenBank database. No BLAST hit matched the complete sequence of ColE1-6kb plasmids, but the majority of hits covered approximately 55 % of the plasmid genome. In most cases, these hits corresponded to mobilization genes. For ColE1-4kb, the closest hit was 99 % identical to the colicigenic plasmid of *Salmonella enterica* serovar Berta (pBERT) and covered the entire genome. All other plasmid hits covered <82 % of the ColE1-4kb genome and were represented in various species of Gram-negative bacteria. The ColpVC-2kb was >99 % identical to plasmids from 5 *Salmonella* sp, including three *S.* Heidelberg strains isolated from Chicken in Canada (data not shown). These results suggested that complete sequences of Col plasmids of PL origin are not well represented in GenBank.

To address this possibility, we performed a BLAST search against the NCBI whole genome shotgun contigs database restricting our search to only serovar *Salmonella*. This database contains genome assemblies of incomplete genomes or incomplete chromosomes of bacteria that were sequenced by the shotgun strategy. *Salmonella* Heidelberg and *S*. Typhimurium strains were the major serovars with contigs matching >99 % of the ColE1-6kb plasmids. For instance, the top 5 contigs were 100 % identical and were 6183 bp long (NCBI Bioproject PRJNA417775). These contigs were detected in S. Heidelberg strains collected from broiler breeder farms in the USA and the isolation source was environmental booties. The ColE1-4kb and ColpVC-2kb plasmid were detected in contigs from sequenced genomes of multiple serovars including Heidelberg, Rissen, Hadar, Typhimurium, Albert, Kentucky, Infantis, Enteritidis etc. The percent identity and query cover for ColE1-4kb and ColpVC-2kb across the serovars ranged from 97 -100 % and 64 -100 %, respectively. Surprisingly, majority of the S. Heidelberg recovered from the broiler breeder farms carried multiple Col plasmids (data not shown). For example, strain C_NS-007 (Accession: SAMN08031253) harbored 3 Col plasmids of sizes ~6 kb, 4 kb and 2 kb encoding the same proteins as observed in this study (Supplementary Fig. 4).

Furthermore, we determined the concentration of Col plasmids in PL collected from conventional and backyard poultry farms located in 4 states in USA. The Col plasmid genes were detected in all litter types except 1 backyard litter, whereas the Rep protein for IncX1 was detected in only 1 of 7 of the litter types, suggesting that Col plasmid genes are abundant in PL (Fig. 7a).

**Figure 7.**
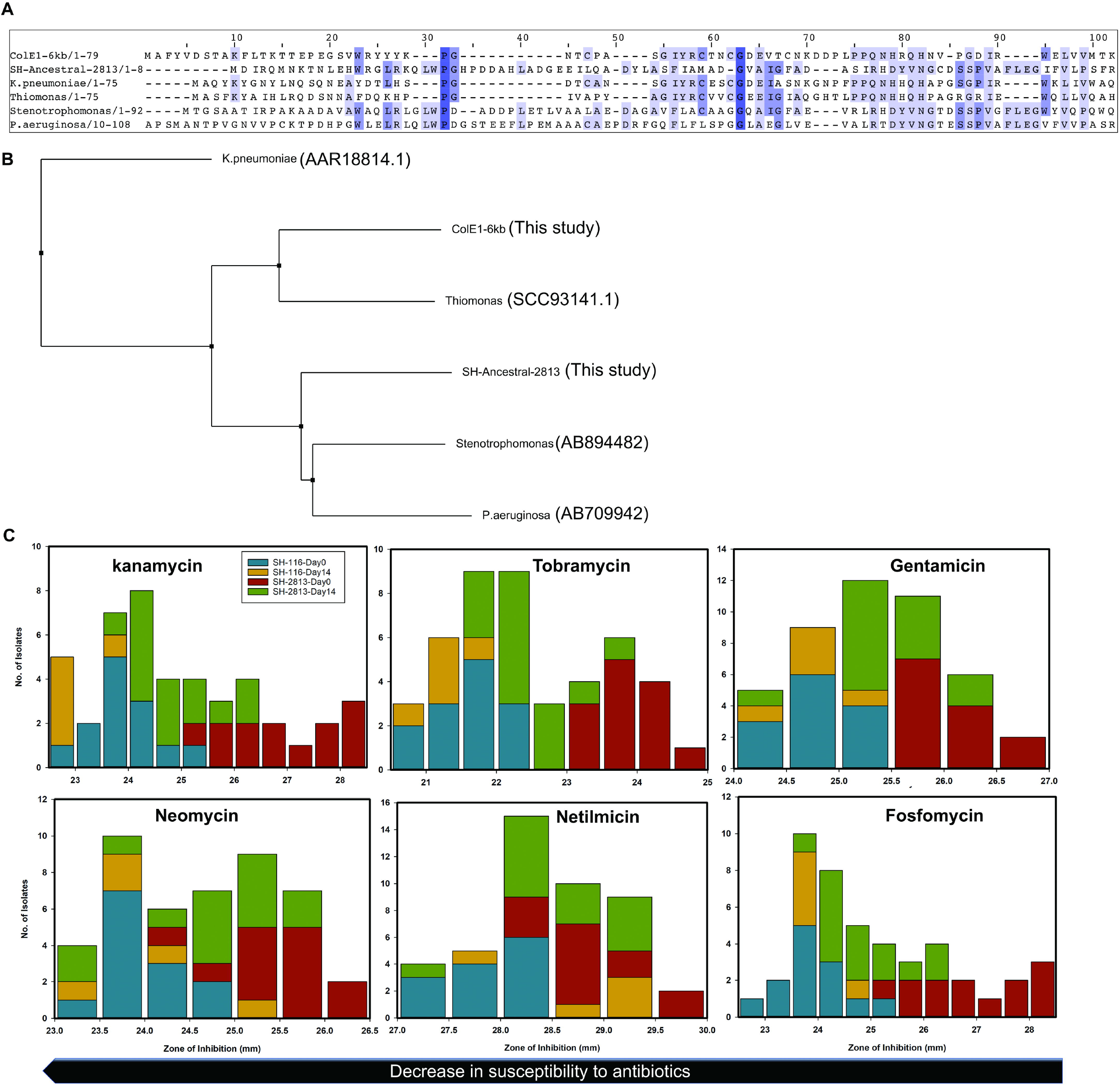
Col plasmids present in poultry litter (PL). **(a)** Concentration of plasmid-specific genes and 16S rDNA in poultry litter of different origins. PL was collected as grab samples from multiple locations of various poultry houses/farms in USA and composited in 1-gallon Whirl-Pak bags. qPCR was performed on DNA extracted from 250 mg of PL in triplicates and pooled. Error bars represent the standard error of the means. Symbols: Conventional farming (Con), Backyard farming (Back), replication protein of IncX1 (IncX1-rep), replication protein of ColpVC (ColpVC-rep), abortive phage infection protein of ColE1-6kb (ColE1-6kb-Abi), macrophage stimulating factor protein of ColE1-4kb (ColE1-4kb-MSF) and 16S ribosomal DNA of Enterobacteriaceae (16S rDNA). **(b)** Concentration of plasmid-specific genes in PL used for microevolution experiment. DNA was extracted from 48 microcosms and qPCR was performed using primers targeting the incRNAI region of ColE1-6kb and ColE1-4kb, the replication protein of ColpVC and the 16S ribosomal DNA of Enterobacteriaceae. Note. One uninoculated microcosm was included at day 0. ColE1 was not detected in SH-2813-BHI and SH-116-BHI microcosms. \**p*.value =0.086 (Wilcoxon signed-rank Test). The error bars represent the standard error of the mean.

### Poultry litter as a selection environment for prophages

Although plasmids are the major MGE elements that have been shown to be rapidly transferred between bacterial populations due to the high number of plasmid copies found per cell — the loss or uptake of prophages cannot be overlooked. Prophages can propagate horizontally between cells (lytic phages) or vertically if they are lysogenized, typically by integrating into bacteria chromosome as prophages. Prophages carry several virulence and accessory genes that can increase their fitness under certain environmental conditions, such as nutrient limitation, oxidative stress, biofilm formation and antibiotic resistance (58). However, intact prophages are like molecular time bombs that kill their hosts upon activation of lytic cycle (59, 60). Consequently, one should expect prophage inactivation to be under strong selection in the environment (59).

To determine if prophage selection played any role during the our microevolution experiments, we used PHAST (24) for predictions of prophages in sequenced genomes. The ancestral S. Heidelberg strains each carried 3 intact (Gifsy-2, PaV-LD, sal-4) and 2 defective prophages (Gifsy-1, BcepMu). Only phages found intact by PHAST were further analyzed in this study. Gifsy-2 (NC_010393) prophage harbored a Cu-Zn superoxide dismutase (SOD) and an attachment invasion locus (OmpX) gene with known oxidative stress (61, 62) and virulence (63) functions respectively. PaV-LD (NC_016564) is a planktonic phage carrying Salmochelins (siderophores of *Salmonella* enterica and uropathogenic *Escherichia coli* strains) transcribed from *iroBCDEN* genes and sal-4 (NC_030919) carried accessory gene for polymyxin resistance (*arnC).* We observed that the sal-4 prophage shared the same proteins as enteroP22 and sal-5, and the number of query hits matching them was similar (data not shown). For convenience, we have named all prophages predicted as P22 or sal-5, sal-4 in this study. The prophage Gifsy-2, PaV-LD and sal-4 were found intact in 84, 19, and 45 of the 85 sequenced evolved strains, respectively. Strain SH-32A-116-Day7-PLE had no intact phage (Supplementary Fig. 5).

## DISCUSSION

### Positive selection for hotspot mutations during microevolution in PL

Repeated, phylogenetically unlinked nucleotide changes that results in mutation of the same amino acid positions are called hotspot mutations (64–66). Two common types of hotspot mutations have been described: (1) parallel mutations that lead to the same amino acid change in the same position and (ii) coincidental mutations that lead to a different amino acid change in the same position (64). These forms of convergent evolution are strong indicators of positive selection and not expected to occur by chance alone (65). We identified 37 parallel mutations carried by all SH-116 isolates recovered and in SH-2813-Day7 and SH-2813BPW-Day1 isolates. A significant proportion of these mutations were point mutations identified in proteins with multiple functions. For instance, a non-synonymous mutation (gln123stop) was present in the cyclic di-GMP regulator (CdgR) — a universal bacterial second messenger that regulates a variety of processes, including cell motility, intercellular interactions, biofilm formation and dispersal and responses to diverse environmental changes (67). The amino acid change resulted in a premature stop codon in position 123. A potential effect of such mutation is a truncated, incomplete or nonfunctional CdgR. High levels of *cdgR* have been demonstrated to promote biofilm formation and sessility, while low levels promote motility and the synthesis of virulence factors in human and animal pathogens (68). A mutation (val196Asn) was also identified in a two-component system sensor histidine kinase (EnvZ) that is involved in the regulation of osmoregulation and controls the expression of outer membrane porin proteins OmpF and OmpC (69). Missense mutation in *envZ* gene has been shown to reduce the expression of porin genes, thus reducing the permeability of the outer membrane of *Salmonella* to cephalosporin antibiotics (70).

Non-synonymous mutations identified in ferric-coprogen receptor (FhuE) and tonB-dependent vitamin B12 (BtuB) suggest that these mutations are related. FhuE serves as a receptor for ferric-coprogen and ferric-rhodotorulic acid and uptake of ferric-coprogen is BtuB dependent. Together, they are required for the uptake and utilization of cobalamins and siderophores (71, 72). These two genes are controlled by Fur regulator and a RNA riboswitch, and their expression is repressed when iron and cyanocobolamin reaches a certain level (71). The effect of this mutation on their interaction or expression in *S*. Heidelberg is beyond the scope of this study. Multiple SNPs/InDels was identified in the 23S-5S ITS region of isolates with hotspot mutations. A large fraction of these mutations were insertions ranging in size from 2 – 70 bp and were found in multiple loci within the ITS region (4035545 – 4035576). The high number and frequency of these mutations may reflect a recent emergence of new lineages and/or their relative evolutionary instability (64). The 23S-5S rRNA ITS region of *Escherichia coli (E. coli)* have been shown to be the most hypervariable region found throughout the ribosomal RNA (rrn) operon, and has also been reported for *Salmonella* Typhi (73). To assess this variability across S. Heidelberg genomes, we reconstructed a phylogenetic tree with the 23S-5S ITS DNA sequences from 22 complete genomes of S. Heidelberg available in GenBank (Supplementary Fig. 6). In addition, we included the ITS region for 14 isolates from this study. The GenBank strains clustered with either day 0 or day 14 isolates from this study. One explanation could be that the strains clustering with day 14 isolates have been co-selected by a recent common environmental pressure such as PL, whereas the strains clustering with day 0 isolates have not undergone a common selection pressure. However, it is important to mention that the GenBank strains (19 of 22 strains) were isolated in Canada and may have biased the result of the reconstructed tree. Another possible explanation is that this region of the *rrn* operon may be prone to indel accumulation and is relatively unstable. For example, mutation was not identified in this region in some of the isolates having hotspot mutations for both SH-2813 (6 isolates) and SH-116 (6 isolates).

High frequency hotspot mutations have been identified in the core genomes of *E. coli* and *Salmonella* serovar *enterica* (64, 66, 74). Furthermore, hotspot mutations are more likely to be of recent origin than long term (65). Chattopadhyay et al (64) performed an in-depth analysis on hotspot mutations in multiple *E. coli*, *Shigella* and *Salmonella* sequence types and concluded that the number of recent hotspot mutations were significantly higher than long term ones. In addition, the frequency of recent hotspot mutations increased by the addition of more genomes to their analysis. Taken together, our result can be explained by the so-called “source sink” dynamics of pathogen microevolution. The “source” can be defined as an evolutionary stable habitat (e.g. chicken gut), while the “sink” as an unstable transient compartment of a habitat (e.g. poultry litter) where S. Heidelberg is introduced occasionally. Under this scenario, hotspot mutations that are adaptive for survival in PL are continuously selected for on a short term, but may be selected out upon colonization or entry into the original stable habitat. This tradeoff may result in different hotspot mutations in *S.* Heidelberg strains isolated from PL, chicken gut or humans.

### Col plasmids and IncX1 contributed to the fitness of *S.* Heidelberg

In this study, only *S.* Heidelberg strains carrying 2 ColE1 and cryptic ColpVC plasmids survived more than 7 days in used PL. The ColE1-6kb relaxases (MbeA) shared >77 % homology with the relaxase of ColE1-4kb but was more closely related to the prototype ColE1 (>96 %) plasmid of *E. coli.* When the ColE1 plasmids sequences from this study were compared to those available in NCBI, we found that these plasmids were underrepresented in the nr database but abundant in the genome assemblies of incompletely assembled *Salmonella* genomes. The observed low frequency of complete ColE1 plasmids harbored by *Salmonella* in NCBI may be a consequence of the bias towards sequencing strains carrying plasmids with known antibiotic resistance genes or virulence. After screening multiple PL of diverse origin with qPCR, we detected Col plasmid genes at high levels in 6 of 7 litter types. Our data further suggest that the PL environment may impose selective pressure that promotes the acquisition of these Col plasmids.

The ColE1-6kb plasmids were present in higher copy numbers than ColE1-4kb and carried 2 putative genes encoding a type-IV TA system and an aminoglycoside-acetyltransferase. The TA system was linked by STRINGDB (75) to the widespread abortive bacterial infection (Abi) system belonging to the cluster of orthologous group COG1106 (domain of ATPases) and the closest homologs were those present in Pseudomonadaceae and Lactobacillaceae. In some bacteria, this protein is annotated as ATP-binding protein or abortive infection protein. Abi systems induce phage resistance to prophages of type 936 in *Lactococcus* and enhance plasmid maintenance/stabilization via bacteriostatic toxicity (76). Consequently, cells carrying plasmids encoding the Abi gene will be immune to phage attack or replication, thereby providing protection for their clonal population. Their role in *Salmonella* remains questionable; however, their high expression under laboratory growth conditions in our study (Supplementary Fig. 7) suggests that they are functional and may have contributed to the persistence of S. Heidelberg in PL.

We do not know the potential functions of the accessory genes on ColE1-4kb, but their compatibility with ColE1-6kb may stem from differences in their RNAI sequences (77) and their mob region. The ColE1 plasmid family represents plasmids that are only transmissible in the presence of an additional plasmid with conjugative functions and usually carry a mob region encoding specific relaxase and accessory mob genes (*mbeB, mbeD mbeC).* The relaxase component is crucial for both initiation and transfer of DNA and is highly conserved (78, 79). In addition, it has been predicted that the C-terminal region of MbeA binds ssDNA, while the N-terminal fragment contacts MbeC (80, 81). The observed differences in C and N-terminal of MbeA and MbeC in this study suggests that such modifications may be critical for successful coexistence of multiple ColE1 plasmids in one host cell. We used these conserved mob protein sequences for phylogenetic tree reconstruction and assessed if there is evolutionary relatedness between the ColE1 plasmids from this study. The majority of the isolates (11 of 14) included in the MbeC tree were from SH-116, while the 3 isolates included in the MbeA tree were from SH-2813 strain. These results infers that the ColE1 plasmids carried by both SH-2813_evol_ and SH-116 isolates have identical relaxases but differ in MbeC proteins.

The Incompatibility factor X1 plasmid harbored by the S. Heidelberg strains used in our study more than likely served a conjugative role. IncX1 is a low copy plasmid and carries required mating pair formation system for horizontal gene transfer (HGT) including type IV secretion systems (T4SS), coupling proteins and a relaxase (82, 83) (Fig. 2a). This plasmid was present in all strains recovered which infers that they are stably maintained and critical to the survival S. Heidelberg strains carrying them. Stability of conjugative plasmids has been attributed to high conjugation rates and plasmid stabilization proteins (84, 85). The InDel present in VirB8 resulted in a frame-shift mutation in this protein and subsequent changes in DNA regions coding for T4SS proteins. Indeed, this mutation resulted in the fusion of a smaller VirB8 (4.4 kDa) encoding gene with a bigger VirB8 gene (23.4 kDa) (Fig. 2a). In addition, differences were observed in sequences coding for the conjugal transfer protein (TraG), an important protein for bacterial conjugation (86). Interestingly, one strain without TraG (SH-33A0.5-116-Day7-PLE) was isolated but we do not know the conjugal implication of this deletion (Supplementary Fig. 8)

Following the acquisition of Col plasmids, we observed a significant reduction in the expression of replication protein (Rep) protein and a type II toxin-antitoxin (TA) gene. This reduction in *rep* expression suggests that *rep* expression may be linked to the cost of taking up new MGE. Reducing *rep* expression will subsequently decrease the copies of IncX1 available for ColE1 plasmids mobilization — thus reducing the burden associated with maintaining multiple plasmids. Also, regression analysis of IncX1 and Col copies showed that IncX1 was a good predictor of ColE1-6kb copies per cell — evolved strains carrying high IncX1 copies also carried high ColE1-6kb copies (Fig. 3; Supplementary Table 6). This result suggests there might be equilibrium between the numbers of “helper plasmids” to mobilizable plasmids residing in a cell.

Bacteria (TA) modules are genetic elements composed of a toxin protein component that disrupts bacterial growth by interfering with vital cell processes and an antitoxin that impairs the functionality of the toxin until this inhibition is abolished in response to cellular signaling. Type II TA system carried on plasmids are important players in post-segregational killing — plasmid-less cells are killed by the toxin, thereby establishing the dominance of plasmid-bearing cells in a population (52, 53, 87). At late logarithm and stationary phase growth in Mueller Hinton Broth, the *relB* gene (antitoxin) of IncX1 was significantly downregulated in evolved strains compared to ancestral strains, and no change was observed in *relE* (toxin) expression (Fig. 2c). This continuous expression of *relE* will ensure that only IncX1 carrying cells are present in the population. Therefore, disrupting the conjugative and stability genes of conjugative plasmids may represent a critical point for limiting their persistence in the environment. For instance, a combination treatment with linoleic and phenothiazine reduced the conjugation efficiency of six conjugative plasmids by 3 -fold, including the IncX plasmid of RP4 (88).

Furthermore, the IncX1 present in Col plasmid-bearing strains of S. Heidelberg carried a 22-bp direct repeat deletion at the origin of replication (*ori),* an Indel in VirB8 (G-> GA) and a non-synonymous substitution in a hypothetical protein (I97T). Isolates with this deletion had higher copies of IncX1 than ancestral strain carrying the extra 22-bp direct repeat. This result supports other studies showing that the deletion of iterons increased plasmid copy number of theta plasmids (reviewed in (90)). For example, Millan et al (91) showed that mutations acquired in the OriV of a synthetic ColE1-like plasmid (pBGT) carrying bla_TEM-1_ β-lactamase gene increased plasmid copy number and subsequently increased ceftazidime resistance several folds.

To expand beyond the *S*. Heidelberg strains used in our study, we performed an alignment of the *ori* of closed “circular” IncX1 plasmids of approximately 37kb present in *S.* Heidelberg strains available in NCBI. We observed that all the strains analyzed carried only two iterons (Fig. 2b), which implies that this mutation is overrepresented in sequenced *S*. Heidelberg carrying IncX1 plasmids. The majority of these strains were from clinical settings (Supplementary Table 8) which suggests they are being recovered more often and may be associated with an increase in fitness of those strains as was also observed in our study. Taken together, our results support the idea that iterons regulate IncX1 plasmid replication and their compatibility with other plasmids.

Mutations acquired on plasmids and chromosomes after an HGT event have been shown to compensate for the fitness cost associated with the acquisition of a new MGE (43, 91–93). Here, strains carrying Col plasmids had a total of 4 plasmids and persisted for up to 14 days, which is a “plasmid paradox,” where the acquisition and maintenance of multiple plasmids over the long term is supposed to impose a burden on their host (84, 85). This principle has been elucidated upon, where positive epistasis minimized the cost associated with carrying small plasmids of *Pseudomonas aeruginosa* over a long-time scale (42). Recently, a study showed that the acquisition of three ColE1 plasmids in naïve *H. influenzae* hosts did not impose a fitness cost as long as the bacterium compensated for the first plasmid (94). The presence of one plasmid favored the presence of a second plasmid and the presence of this second plasmid or third plasmid did not impose an additional significant biological cost, even though these plasmids produced a cost when they are alone in the cell. Col plasmid-bearing isolates from this study had increased fitness compared to a isolate without a Col plasmid and carried 37 hotspot mutations (Fig. 1d). It is unlikely that all observed SNPs/InDels are to compensate for Col plasmid acquisition, as some will be compensatory for other MGE such as bacteriophages (95–97). For example, the prophage Gifsy-2 was found intact in 97% of S. Heidelberg isolates sequenced in our study, suggesting the PL environment may have selected them for increased fitness (Supplementary Fig. 5). Further, the ubiquity of Gifsy-2 in S. Heidelberg strains has been reported, which infers they are important for S. Heidelberg survival (13). We identified a hotspot mutation in the intergenic region upstream of cell division protein (FtsZ) of Gifsy-2, an important protein for cell division and lambda bacteriophage infection in *E. coli* (98) (Table 2). In summary, it is more than likely that the hotspot mutations observed in this study are associated with HGT events involving plasmids and prophages, in addition to mutations selected for persistence in PL.

### Plasmid and chromosome encoded genes mediated antibiotic resistance in *S*. Heidelberg

Another significant observation from this study is the decrease in susceptibility of Col plasmid-bearing strains towards kanamycin, tobramycin and fosfomycin. We do not know what these reductions in inhibition zones mean in terms of MIC as there are no current CLSI standards (46) for *S*. Heidelberg. In addition, it has been shown that simple back-calculations from inhibition zones to MIC can be confounded by several covariates such as serotype, host species, and year of isolation (99). The chickens raised on the PL used for this study were not administered any drugs (Personal Communication-Dr. Casey Ritz, University of Georgia), thus experimental selection for these resistance phenotypes is highly unlikely. It is a possibility that Col plasmid selection or population factors like competitive exclusion and the presence of “cheaters” (100–102) could account for this result but more research will be needed to test these possibilities.

Despite the fact that the acquisition of Col plasmids coincided with the observed change in susceptibility, we cannot say for sure if these plasmids played any role. The putative aminoglycoside acetyltransferase (aac-(6’)) gene carried on ColE1-6kb represents a fragmented aac-6’ gene and shares significant homology with the N-terminal of aac(6’)-Ib of *K. pneumoniae.* Its small size (81 amino acid residues) compared to other known aac genes (Fig. 5a), and the absence of homologs in all antibiotic resistance genes (ARG) databases queried suggests it may be an uncharacterized ARG. Importantly, its presence on ColE1 will result in more copies of these gene compared to the aac-(6’) gene carried as a single copy on the chromosome. More studies will be required to determine if these two aac-(6’) genes confer decreased susceptibility towards the aforementioned aminoglycosides.

Regarding fosfomycin, Rehmann et al (54) determined that S. Heidelberg strains with an inhibition zone of 26 mm were resistant to >32 μg/ml of fosfomycin, which suggests that evolved *S*. Heidelberg strains carrying Col plasmids in our study may be resistant to fosfomycin. The fosA7 gene is present in many S. Heidelberg genomes and resistance to fosfomycin has been demonstrated in a few (54). In addition, S. Heidelberg harbors other chromosomal genes associated with fosfomycin resistance including *glpT* and *uhpT* (103). We did not observe any mutation in these genes, thus cannot correlate fosfomycin resistance to a specific ARG. This result shows the intrinsic bias that could be associated with estimating resistance phenotype from only known ARG that are available in curated databases. More concerning, is the spontaneous mutant (SH-17BBPW-2813) that emerged with reduced susceptibility to 3 classes of antibiotics following overnight enrichment in buffered peptone water (BPW). One explanation is that BPW enriched for the limited strains present in PL or it created an environment where only the fittest strain emerged. Therefore, we argue that chromosomal mutations in addition to Col plasmids are responsible for the reduced susceptibility displayed by evolved *S*. Heidelberg isolates.

Results from this PL microevolution experiment suggest that chromosomal mutations and Col plasmids contributed to the fitness of *S*. Heidelberg. Col plasmids were present in all *S*. Heidelberg isolates sequenced after 7 days in PL suggesting that their presence is required for survival. However, the population of SH-116 strains were significantly lower in PL at day 14 compared to SH-2813 populations. We expected SH-116 to be in higher abundance than SH-2813 since the ancestral populations carried Col plasmids. This was not the case in this study (Fig. 1a). SH-116 strains were only detected in 1 of 6 microcosms by direct count and in 3 of 6 after enrichment in buffered peptone water (BPW) after 14 days incubation in PL. In contrast, SH-2813 was recovered by direct count in all 6 microcosms (Supplementary Table 1). The dynamics observed in our experiment can be modeled according to Watve et al (104) Rock-Paper-Scissors (RPS) model of sociobiological control of plasmid copy number. In this model, when a mutant bacterium with higher copy number plasmid was introduced in the presence of a wildtype low copy plasmid, the wildtype was driven to extinction. This high copy mutant was then vulnerable to invasion by another mutant that had higher copy numbers, which in turn was vulnerable to invasion by another mutant with higher plasmid copies. In our scenario, SH-116_anc_ population with lower copy ColE1 plasmids were displaced by those with higher copy ColE1 plasmids mobilized from PL microbiota until the point at which the intrinsic survival rate of *S*. Heidelberg was greatly reduced (Fig. 1a). In contrast, a stable co-existence of SH-2813 of low, intermediate and high copy ColE1 plasmids was observed in the SH-2813 population. Based on the RPS model, it is likely that SH-2813 isolates with higher copy plasmids will end up invading isolates with lower copies. Therefore we should expect a system without a unique equilibrium.

The ColE1 plasmids carried by SH-116 isolates had mob protein (MbeC) sequences that were represented by 11 evolved isolates on the reconstructed phylogenetic tree (Fig. 4c), whereas the SH-2813 isolates had only 3 evolved isolates. This suggests that SH-116 may be acquiring new ColE1 plasmids from bacteria present in PL. This observation was corroborated by qPCR on extracted DNA from PL with Col plasmid gene primer sets. At day 0 and 14, the copies of ColE1-6kb and ColpVC were significantly higher in SH-116-PLE microcosms than SH-2813-PLE (Fig. 7b), which supports the notion that the original SH-116 strains may have been displaced by those carrying higher copy ColE1 plasmids. For the BHI microcosms, we did not detect any plasmid gene even though 16S rRNA result showed an abundance of bacterial DNA (data not shown). We have attributed this discrepancy to the significant differences in moisture and pH between BHI and PLE microcosms. To avoid contamination and mix up of strains, evolution experiments with SH-2813 and SH-116 strains were done on different days. In addition, incubation of microcosms was done in separate incubators (similar make and model) and in separate laboratories. The plastic containers with BHI and microcosms were placed on the upper and bottom layer of the incubators. To avoid drying out of PL in incubator, a 2-liter bucket of water was placed on the upper layer next to the BHI microcosms. An unforeseen outcome of this experimental setup was an increase in moisture for BHI microcosms. Moisture has been shown to be a significant factor changing the microbiota of PL (7). Thus, it is plausible that this difference in moisture reduced the population of resident microbiota carrying Col plasmids in the BHI microcosms compared to PLE. Further research from our laboratory will be exploring this hypothesis.

## CONCLUSION

Hotspot mutations and ColE1 plasmids acquired under positive selection from PL contributed to the persistence of *S.* Heidelberg observed in this study; however the identity of the bacterial donors of these Col plasmids remains unknown. Further, due to space limitation we did not discuss in detail exactly how each mutation observed in evolved isolates of S. Heidelberg can confer a fitness advantage in PL. Also, we did not identify mutations that could have been due to homologous intragenic recombination events. Owing to the large amount of information obtained from the SNPs/InDels analyzed across 86 genomes, it was not practical to perform an in-depth analysis on every one of them. Therefore, we have provided a list of all the identified mutations, their corresponding VCF files and UNIX command line used (Supplementary Table 4 and Dryad Digital Repository: https://doi.org/10.5061/dryad.pc6tp4q).

## ACKNOWLEDGEMENTS

We are grateful to Dr. Casey Ritz for providing us with the poultry litter used in this study and for Caroline Plymel for her help with sample analysis. We wish to thank Dr. Karamat Sistani for valuable assistance with litter nutrient analysis. We also appreciate Steven Lankin and Mary Katherine Crews for manuscript review. Any opinions expressed in this paper are those of the authors and do not necessarily reflect the official positions and policies of the USDA and any mention of products or trade names does not constitute recommendation for use. The authors declare no competing commercial interests in relation to the submitted work.

## Supplementary Figures

**Figure S1**. Experimental design.

**Figure S2**. Principal component analysis (PCA) of poultry litter characterizing the differences between microcosms. Data represent 48 microcosms. Positively correlated variables point to the same side of the plot and negative correlated variables point to opposite sides of the graph. Symbol: Colored by contributions to the principal component (Contrib).

**Figure S3**. ProgressiveMauve alignment of (**a**) IncX1 (**b**) ColE1-6kb and (**c**) ColE1-4kb whole plasmid DNA sequence from selected isolates from this study. DNA regions that differ are highlighted with horizontal black rectangular boxes.

**Figure S4**. Linear maps of Col plasmids detected in an incomplete assembly of a S. Heidelberg strain. Strain NS-007 (NCBI accession: SAMN08031253) was isolated from a broiler breeder farm in the USA under Bioproject PRJNA417775. Contig size for ColE1-6kb, ColE1-4kb and ColpVC-2kb are 6183, 4778 and 2105 bp, respectively.

**Figure S5**. Intact prophages in sequenced genomes of S. Heidelberg from this study.

**Figure S6**. Maximum likelihood tree of S. Heidelberg strains available in GenBank based on 23S-5S internal transcribed spacer. The consensus 23S-5S ITS sequence of isolates included from this study were extracted by mapping whole genome sequence reads to *S*. Heidelberg reference genome (CP016573)

**Figure S7**. Gene expression of selected Col plasmid genes determined by qRT-PCR for (**a**) SH-116_anc_ (**b**) SH-116_evol_ and (**c**) SH-2813_evol_ isolates. Log2 fold-change is relative to the expression of chromosome encoded reference genes *gapA* and *gyrB.* Symbols: replication protein of ColpVC (ColpVC-rep), abortive phage infection protein of ColE1-6kb (ColE1-6kb-Abi), macrophage stimulating factor protein of ColE1-4kb (ColE1-4kb-MSF) and hypothetical protein (hypo).

**Figure S8**. Circular map of IncX1 plasmid of isolate SH-33A0.5-116-Day7-PLE.

## Supplementary Tables

**Table S1**. Bacteria concentration in poultry litter microcosms.

**Table S2**. Physicochemical properties of poultry litter microcosms.

**Table S3**. Metadata on isolates sequenced.

**Table S4**. SNPs/InDels identified in sequenced S. Heidelberg genomes from this study compared to the reference genome.

**Table S5**. Identified plasmids topology.

**Table S6**. Plasmid copy number determined by whole genome sequencing coverage.

**Table S7**. Antibiotic susceptibility test results.

**Table S8**. Metadata on NCBI S. Heidelberg strains carrying IncX1 plasmid.

**Table S9**. Primers used in this study.

## Supplementary Files

**Supplementary File 1**. Quality assessment and coverage of sequenced SH-2813 strains.

**Supplementary File 2**. Quality assessment and coverage of sequenced SH-116 strains.

**Supplementary File 3**. Disc diffusion inhibition zone measurements.

